# A Cell-Permeable Fluorescent Probe Reveals Temporally Diverse PI(4,5)P2 Dynamics Evoked by Distinct GPCR Agonists in Neurons

**DOI:** 10.1101/2024.06.17.599302

**Authors:** Rajasree Kundu, Samsuzzoha Mondal, Akshay Kapadia, Antara A. Banerjee, Oleksandr A. Kucherak, Andrey S. Klymchenko, Sandhya P. Koushika, Ravindra Venkatramani, Vidita A. Vaidya, Ankona Datta

## Abstract

Lipids, key constituents of cell-membranes, are the first responders to cell signals. At the crux of spatiotemporal dynamics of lipid-signaling response are phosphoinositides. Indeed, phosphoinositides like phosphatidylinositol-(4,5)-bisphosphate (PI(4,5)P2), present in the inner-leaflet of eukaryotic cell-membranes, form the link between signal reception and downstream signal-transmission. In this backdrop, reversible fluorescent probes that can track live PI(4,5)P2 dynamics on a seconds time-scale will afford key insights into lipid-mediated signaling. However, realizing cell-permeable PI(4,5)P2-selective sensors for imaging dynamics remains a challenge due to the presence of structurally similar lipids and low levels of PI(4,5)P2. We report a computationally-designed, rapid-response, reversible, photo-stable, fluorescent sensor that permeates living cells, neurons, and a multicellular organism within few min of direct incubation and distinctly visualizes PI(4,5)P2 pools. We used the sensor to interrogate the role of PI(4,5)P2 in driving heterogeneity of signaling responses and contrasting behavioral effects that ensue upon binding of distinct ligands to the same G protein-coupled receptor. Specifically, we asked whether probing PI(4,5)P2 dynamics using our novel sensor could uncover the earliest of signaling differences evoked by hallucinogenic versus non-hallucinogenic ligands at the serotonin2A (5-HT_2A_) receptor. Our results reveal that a hallucinogenic ligand at the 5-HT_2A_ receptor leads to a slower rate of PI(4,5)P2-depletion when compared to a non-hallucinogenic ligand, within the initial seconds of ligand addition, but has a sustained effect. The ability of our designer chemical probe in timing early seconds-minute timescale lipid-dynamics in living cells opens avenues for tracking early time-point molecular events in neuronal response to chemical and physical stimuli.

## Main Text

The plasma membrane is far more than a mere structural boundary, it is the interface where signal transduction commences, a gateway for all cellular communication, and an active signaling platform.^1^ Central to the roles of the plasma membrane in cellular processes are phospholipids (PLs) that make the membrane.^2^ The localization/distribution/level of a phospholipid in a cell membrane is highly dynamic and changes temporally in seconds to minutes.^3,4^ Tracking this dynamic change in phospholipid distribution and levels in a cell membrane, live, is necessary to understand how the membrane regulates and times cellular signals, transport, and communication.

In the playground of lipid-mediated signal transmission and cellular regulation, phosphoinositides are singularly important.^4–7^ Phosphoinositides, generated by reversible phosphorylation of an inositol-headgroup (**Fig. S1**), are almost exclusively present in the inner leaflet of cell membranes toward the cytoplasmic interface, and in organelle membranes.^4^ The levels of phosphoinositides in cell membranes are ∼1-5% of the total phospholipid content of the membrane.^4,8^ While low in abundance, these lipids are functionally critical and not only mediate classical signal transduction like neurotransmission via G protein-coupled receptors (GPCRs),^7^ but also regulate ion channels,^9^ and impinge on key cellular processes such as exocytosis and endocytosis.^4,5^ Recent studies reveal that both the composition and localization of specific phosphoinositides in proximity to membrane proteins such as GPCRs can influence GPCR activity.^10^ Hence, by visualizing phosphoinositide dynamics within cell membranes, we can gain a direct handle into elucidating the link between lipid dynamics and cellular signaling processes. We asked if distinct ligands at the same GPCR, that ultimately drive divergent behavioral effects, show phosphoinositide-mediated signaling differences as early as few seconds into signal initiation.

A major challenge that we needed to overcome in order to address this question was the ability to selectively visualize the seconds-minute dynamics of a key signal-mediating phosphoinositide, phosphatidylinositol-(4,5)-bisphosphate (PI(4,5)P2), within living cells and neurons. Why is that so? The current gold-standard in PI(4,5)P2 imaging relies heavily on non-responsive protein-based fluorescent sensors (**Table-S1**).^11^ These large probes are diffusion-limited^12^ and especially in the crowded intracellular environment often cannot catch subtle differences in lipid dynamics in seconds-minute timescale,^3,13^ specifically, apparently small but functionally significant effects related to neurotransmission. An example is the differences in early time point signaling events that occur when distinct ligands bind the same cell-membrane receptor.^14^ Apart from their large size, these probes, based on protein-domains that bind to PI(4,5)P2, often cannot distinguish between PI(4,5)P2 and its soluble headgroup counterpart which is a second messenger. Fluorescently tagged artificial phosphoinositides are an alternative to protein-based sensors.^15–17^ However, modified lipids can be mis-localized in the cell membrane due to their altered chemical properties compared to native lipids.^6,16,18^ Importantly, their in-cell dynamics is different from native counterparts.^18^

Here, we report that a rapid response, reversible, ratiometric, cell-permeable short (12 amino acid) peptide-based, photo-stable, fluorescent, PI(4,5)P2-selective sensor, times the early seconds-minute differences in signaling events at the cell membrane of neurons in response to the binding of a hallucinogenic versus a non-hallucinogenic agonist of the serotonin_2A_ (5-HT_2A_) receptor. These results open avenues to access the earliest of events in lipid-mediated signaling that can in turn drive differential signaling outcomes, using cell-permeable lipid responsive chemical tools. Finally, the novel sensor, not only tracked PI(4,5)P2 selectively in living cells and neurons but could also be applied in a multicellular model system to visualize the distribution of the lipid.

## Designing a short-peptide based, ratiometric, rapid-response, cell-permeable, fluorescent probe for PI(4,5)P2

First, we set-forth to design and develop a low-molecular weight (1-3 kDa), cell-permeable, rapid-response, photo-stable, fluorescent probe for PI(4,5)P2. PI(4,5)P2 (**Fig. S1**) regulates classical signal transduction. When a GPCR binds to a chemical ligand like a neurotransmitter or an agonist, the enzyme phospholipase C (PLC) cleaves the headgroup of PI(4,5)P2, to release inositol-1,4,5-trisphosphate (IP3) which then acts as a second messenger in signaling pathways (**Fig. S10**).^19^ PI(4,5)P2 is almost exclusively present in the inner leaflet of the plasma membrane. Since we wanted to track and time initial PI(4,5)P2-mediated signaling events during ligand/agonist-receptor interaction at the cell-membrane, we needed a cell-permeable probe that would selectively and sensitively respond to PI(4,5)P2 over other cellular PLs.

A powerful approach in the design of probes is to use polarity-sensitive dyes, which change their emission color in response to local microenvironment variations produced by bio-molecular interactions.^20^ In our previous work we had developed peptide-based fluorescent, ratiometric probes for phosphoinositides which responded to both PI(4,5)P2 and phosphatidylinositol-4-phosphate (PI(4)P) (**Fig. S1**) without considerable specificity toward either phosphoinositide which have similar relative cellular abundances.^8,21^ The probe design involved attaching a polarity sensitive dansyl dye to a phosphoinositide binding peptide segment of the actin-regulatory protein Gelsolin (Gel 150-169). These phosphoinositide probes had low photo- stability which precluded applications in imaging live cell dynamics over several min. Therefore, key features that we desired in a PI(4,5)P2 probe that would track live PI(4,5)P2 dynamics were, selectivity, rapid cell-permeability in min timescale for easy applicability, sensitivity toward lower ∼ 1 mol% PI(4,5)P2, reversible response to visualize dynamics, and importantly photo-stability for time-lapse imaging over several minutes. Our strategy for imaging PI(4,5)P2 dynamics was to design a probe with a cell-permeable peptide-binding unit that would bind to PI(4,5)P2 and provide a specific conformational change that could be leveraged to sense the lipid selectively. The workflow that we devised to attain a PI(4,5)P2 selective probe/sensor involved molecular dynamics simulations on Gel 150-169 (Gel-20aa) and Gel 158-169 (Gel-12aa) peptides in the presence of different phosphoinositide headgroups. The driver for the selection of these peptides was their proven ability to bind to phosphoinositides even when removed from the parent protein.^21^ We asked if there were any specific differences in the interaction of the peptides with different phosphoinositides, that could be leveraged to generate a PI(4,5)P2 selective probe.

All atom molecular dynamics (MD) simulations were performed on both the Gel-20aa and Gel-12aa peptides on a phospholipid bilayer (**Table S2**). The Gel-20aa peptide was reported to adopt a helical conformation in the presence of PI(4,5)P2, hence, an NMR structure of the helical form (PDB ID: 1SOL)^22^ of the peptide was used as a model for the simulations. The structure of the shorter Gel-12aa sequence was modeled based on the structure of Gel-20aa by deleting the eight C-terminal amino acids from the Gel-20aa sequence *in silico*. Three 100 Å x 100 Å membrane bilayers made of phosphatidylcholine (PC) molecules were created where a single phosphoinositide molecule, either PI(4)P or PI(4,5)P2 or phosphatidylinositol-3,4,5- trisphosphate, (PI(3,4,5)P3) (**Fig. S1**), was inserted into one leaflet of the PC membrane to afford 0.74 mol% of PI(4,5)P2 on a single leaflet of the bilayer (**Fig. 1a-c**). This value was within the expected range of the biological abundance (0.5-1 mol%) reported for PI(4,5)P2 and PI(4)P.^8,23^ PI(3,4,5)P3 while at least 100-1000 fold lower in abundance than PI(4,5)P2 and PI(4)P,^8^ was included in the calculations as a control because of its high-structural similarity to PI(4,5)P2. In our simulations, the positively charged residues in the C-terminus of Gel- 20aa were seen to form electrostatic contacts with the negatively charged phosphate moieties of the phosphoinositide headgroups (**Fig. 1a-c, Video-1**). The binding modes of the peptide C-terminus with each of the three phosphoinositides were different. For example, R20 and K17 formed electrostatic contacts with PI(4,5)P2 throughout the entire course of the simulations (**Fig. 1b, Video-1**), while R20 and R19 formed electrostatic contacts with PI(3,4,5)P3 (**Fig. 1c, Video-1**). We estimated average non-bonding interaction energies (***ΔE_NB_***) between positively charged residues of the peptide lying within 3 Å of a phosphoinositide headgroup, and the phosphoinositide headgroup. The peptide showed the highest ***ΔE_NB_*** toward PI(3,4,5)P3 followed by PI(4,5)P2 and PI(4)P (**Fig. 1d, Fig. S4b, Table S3**). Over the course of the 200 ns simulations, within the first ∼13 ns, the peptide aligned parallelly to the membrane allowing other residues as well as the N-terminus to interact with the membrane (**Video-1**). The N-terminus of the peptide dipped into the membrane multiple times when PI(4,5)P2 was present, within the first 30 ns of the simulation (**Fig. S4a**). The reproducibility of the event was confirmed by running additional short 30 ns simulations (**Fig. S4a**). We noted that the N-terminus inserted itself deeper into the membrane for PI(4,5)P2 simulations when compared to that of PI(4)P and PI(3,4,5)P3 (**Fig. 1e, Table S3**). The percentage helicity for the Gel-20aa peptide was the lowest for the simulation performed in the presence of PI(4,5)P2 (**Fig. 1f**), throughout the course of the simulation.

**Fig 1.**
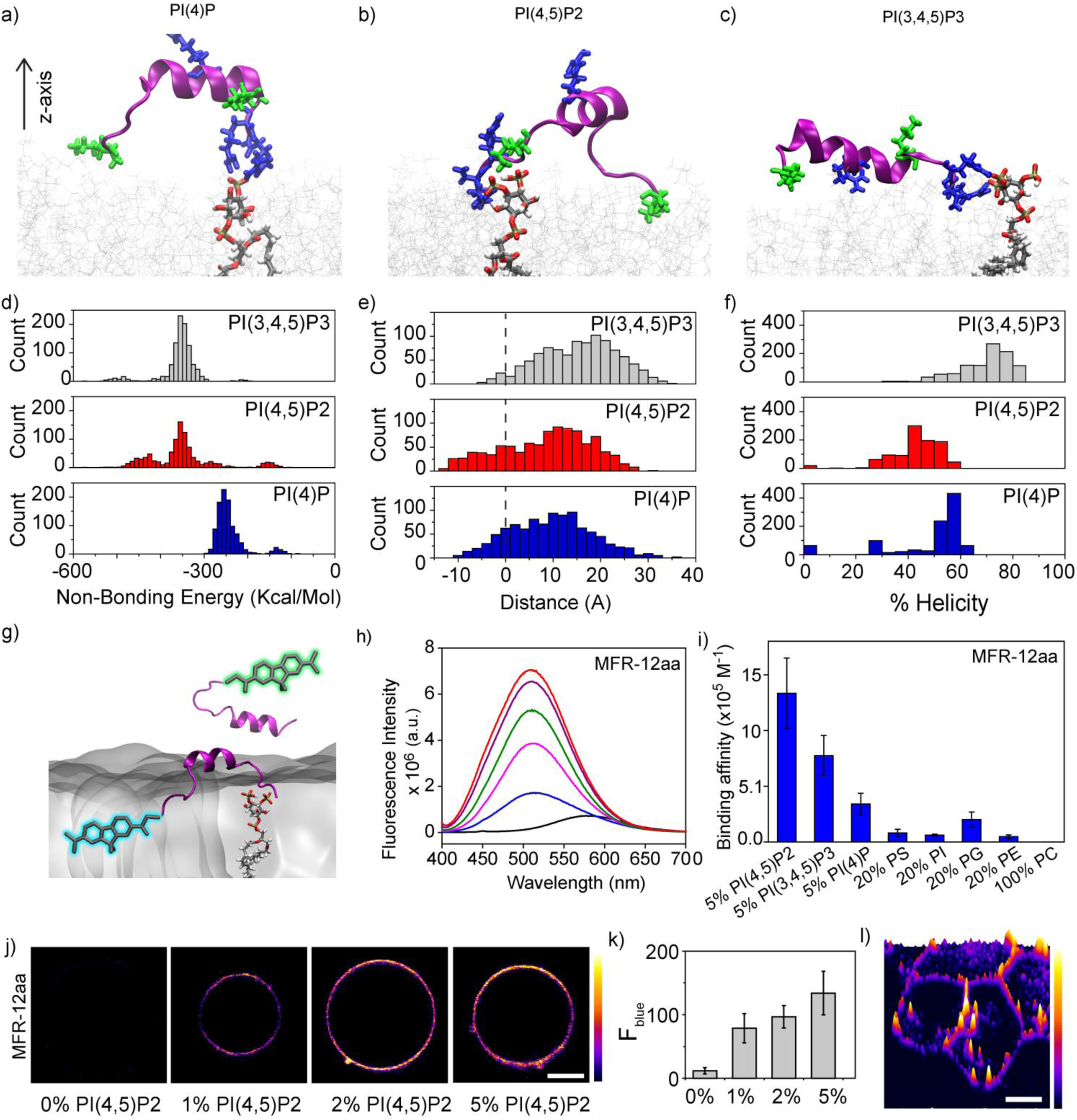
A peptide-based ratiometric fluorescent probe detects PI(4,5)P2 selectively. Snapshots from MD simulations showing Gel-20aa-phosphoinositide interaction (phosphoinositide gray licorice, O-atoms in red) in a PC (gray lines) membrane bilayer: (a) PI(4)P, (b) PI(4,5)P2, and (c) PI(3,4,5)P3. The peptide backbone is in magenta and cationic amino acids are shown in blue (Arg) and green (Lys). (d) Distributions of non-bonding energies for binding of the Gel-20aa peptide to phosphoinositide headgroups. (e) Distributions of the distance between the N-terminus amine group of Gel-20aa and the membrane surface (dashed line) along the z-axis as shown in Fig. 1a. (f) Distributions of helicity content of the Gel-20aa peptide upon binding to phosphoinositide headgroups. All distributions shown in d-f have been calculated from 200 ns MD simulation trajectories. (g) Schematic depiction of sensing strategy used for PI(4,5)P2 detection. (h) Fluorescence response of **MFR-12aa** (1 μM) depicting a blue shift in emission maxima with increasing concentrations (0, 5, 10, 25, 71, and 114 µM) of 5% PI(4,5)P2-PC SUVs, λ_ex_ 390 nm. (i) Bar plot (mean ± SEM., *N* = 3) comparing the association constants (K_a_) of **MFR-12aa** toward various PLs, derived from fluorescence titrations. (j) Confocal images of GUVs containing PI(4,5)P2 (x% where x = 0, 1, 2, 5) in PS (20%)-PC (80-x%) incubated with **MFR-12aa** (200 nM). (k) Bar plots representing average intensity along the perimeter of GUVs obtained from calculations on 10 vesicles. Error bars represent SEM, *N*=3. (l) 3D intensity plot generated from confocal microscopy image of living HEK293T cells incubated with **MFR-12aa** (5 μM) for 15 min at room temperature. Confocal images (j, l) were recorded in blue channel (480-520 nm, λ_ex_ 780 nm) and are representative of 3 independent experiments. Scale bars, 10 µm.

Based on these results we concluded that the N-terminus of the Gel-20aa peptide afforded differential interaction with the membrane in the presence of different phosphoinositides. A closer inspection of the structures of the peptide on membranes with the three different phosphoinositides, hinted that whenever the residue K17 formed an electrostatic contact with the headgroup, the helicity of the peptide was lower (**Video- 1, Fig. 1f**). Specifically, in the case of the interaction of the peptide with PI(3,4,5)P3 where K17 did not make any contact with the headgroup throughout the course of the 200 ns simulation, the percentage helicity was higher than the other phosphoinositides (**Video-1, Fig. 1f**). The conformational flexibility arising from the opening of the helix when K17 interacted with the headgroup might lead to increased membrane dipping of the Gel-20aa N-terminus observed for PI(4,5)P2.

The interaction of the shorter peptide, Gel-12aa, with different phosphoinositide headgroups (**Fig. S3a-c**) was similar to that of Gel-20aa (**Video-2**). This was not unexpected because the C-termini of both the peptides were identical. The non-bonding energies of the Gel-12aa peptide toward different phosphoinositide headgroups followed a similar trend as Gel-20aa (**Fig. S3d-f)**, except that the ***ΔE_NB_*** values of the peptide interacting with PI(3,4,5)P3 and PI(4,5)P2 were now closer, **(Fig. S4c, Table S3**). We noted that despite the differences in N-terminus residues of both peptides, the N-terminus dipping feature was retained over the course of all simulations for the shorter peptide (**Video-2, Fig. S3g-i**) with the maximum dipping observed for PI(4,5)P2 (**Table S3**). The short peptide was not helical based on the simulations, and hence the differential N-terminal dipping observed for the shorter peptide may be attributed to differences in the electrostatic interactions between the cationic amino acids at the C-terminus and phosphoinositide headgroups.

Taken together, the MD simulations for both the Gel-20aa and the Gel-12aa peptides indicated that the extent of dipping of the N-terminus into the membrane was different for the three phosphoinositides studied. We decided to use this feature to develop PI(4,5)P2 selective fluorescent probes. Since the simulations predicted that the N-termini of the peptides afforded maximum dipping upon interaction with PI(4,5)P2, our strategy was to attach a polarity sensitive dye to the N-termini of both peptides with the expectation that the resultant N-terminal dye-conjugated peptides might act as PI(4,5)P2 probes or sensors (**Fig. 1g**). Next, we needed to select a photo-stable polarity sensitive dye with a significant two-photon absorption cross-section. Photo-stability would allow quantitative imaging of live lipid dynamics and multi- photon excitation at near-infrared^24^ would improve both depth penetrability in tissue samples and the detection sensitivity due to minimal background from autofluorescence. The dye that we zeroed in on was a methyl substituted fluorene dye MFR. Previously, a push-pull fluorene dye FR_0_ was reported,^25^ which exhibited superior two-photon cross-section (σ_2_ 400 GM), sensitivity to polarity, and photo-stability (τ_p_ 360 min in ethanol), compared to popular prodan- and dansyl-based polarity-sensitive dyes.^20^ Here, we applied its dimethylamino-analogue MFR, bearing a carboxylate group, for developing our probes. Two probes, MFR-VVQRLFQVKGRR (**MFR-12aa**) and MFR-KHVVPNEVVVQRLFQVKGRR (**MFR-20aa**) were therefore designed as depicted in **Fig. S2**.

## MFR-20aa and MFR-12aa are PI(4,5)P2 selective and sensitive probes

**MFR-20aa** and **MFR-12aa** were synthesized via a solid-phase synthetic strategy and purified (**Fig S16, S17**). In order to test whether the probes afforded a selective response toward a specific phosphoinositide over other physiologically relevant phospholipids, small unilamellar vesicles (SUVs) of ∼ 100 nm diameter were used as model membranes. SUVs containing 5% phosphoinositide in phosphatidylcholine (PC), the major phospholipid component of eukaryotic cell membranes, were prepared and characterized. SUVs containing 20% of several physiologically more abundant phospholipids including phosphatidylserine (PS), phosphatidylinositol (PI), phosphatidylglycerol (PG), and phosphatidylethanolamine (PE) in PC were also included in the experiments. 100% PC SUVs served as control. Since both the probes were completely water-soluble, all in vitro experiments described henceforth were performed in aqueous buffer.

Upon titrating with 5% PI(4,5)P2 in PC SUVs, a 90 nm blue shift (λ_max_ shifted from 575 nm to 485 nm, λ_ex_ 390 nm) was observed for **MFR-20aa** (**Fig. S5a**). Concomitantly, a 70 times enhancement was observed for the emission intensity at 480 nm, at saturating lipid concentration. Similarly, **MFR-12aa** afforded a 70 nm spectral shift (λ_max_ shifted from 580 nm to 510 nm, λ_ex_ 390 nm) and a 100 times fluorescence enhancement at 480 nm, at saturating lipid concentration (**Fig. 1h**). Hence, **MFR-20aa** and **MFR-12aa** provided dual ratiometric and ‘turn-on’ emission response toward PI(4,5)P2. Further, this spectral response suggests that after the peptide binds to PI(4,5)P2, the MFR dye is transferred from polar aqueous environment to the low-polar environment of the lipid membrane as predicted from our computational studies.

The in vitro fluorescence titration data was fitted to a 1:1 probe: lipid binding model (SI section 8) to obtain association constant (K_a_) values for each probe toward all tested physiologically relevant phospholipids (**Fig. 1i** and **Fig. S5b**). Both probes showed the highest association constants toward PI(4,5)P2. Importantly, the binding toward PI(4)P was significantly lower for both probes when compared to PI(4,5)P2, as predicted by the computational analysis of the MD simulations. **MFR-12aa** and **MFR-20aa** afforded at least 20 times higher affinity toward PI(4,5)P2 over phosphatidylserine (PS) which is the major anionic PL (10-15%) present on the inner leaflet of the eukaryotic plasma membrane (**Fig. 1i** and **Fig. S5b**). Both probes afforded a ∼ 1.7-1.8 times higher affinity toward PI(4,5)P2 over PI(3,4,5)P3. Since cellular PI(3,4,5)P3 levels are 100-1000 times lower than Pi(4,5)P2 levels,^8^ we concluded, that the MFR-labelled peptides could act as selective and sensitive PI(4,5)P2 probes in cellulo. The limit-of-detection (LOD) of **MFR-12aa** toward PI(4,5)P2 was obtained from the in vitro ‘turn-on’ response as 220 nM while that for **MFR-20aa** was obtained as 350 nM. From these results we concluded that the shorter peptide-based probe **MFR-12aa** had a similar selectivity, but higher sensitivity as compared to the longer peptide-based probe **MFR-20aa** and hence would be apt for detection of physiological PI(4,5)P2 levels and visualizing dynamics. Further, the response of **MFR-12aa** toward PI(4,5)P2 remained unaffected in the presence of the soluble second messenger IP3 (**Fig. S6)**. This was an advantage over protein-based sensors which often bind to both PI(4,5)P2 and IP3 leading to low specificity toward PI(4,5)P2. Because the shorter size of the **MFR-12aa** sensor could lead to enhanced cell-permeability, we decided to take it forward to test its applicability toward imaging PI(4,5)P2 dynamics in the specific context of agonist-induced GPCR signaling.

Before proceeding to in cell evaluation of **MFR-12aa**, we further tested the sensitivity of the probe toward detection and imaging of low ∼ 1-2 mol% PI(4,5)P2 in model membranes in a confocal fluorescence microscopy setup. Giant unilamellar vesicles (GUVs) with diameter 20-30 µm were prepared with 0-5 mol% of PI(4,5)P2 in PS (20 mol%) and PC (80-75 mol%). PS was used as a component of the GUVs since it is the most abundant anionic PL in eukaryotic cell membranes.^26^ GUV images were recorded via two-photon excitation at 780 nm. Emission was collected in both the blue channel, 480-520 nm, which represented the emission from mainly the PI(4,5)P2 bound probe and the orange channel, 580-620 nm, which represented emission from both PI(4,5)P2 bound and unbound probe. The **MFR-12aa** probe clearly lighted up GUVs with 1 mol% PI(4,5)P2 and showed a progressive increase in blue/bound channel emission intensity as the mol% of PI(4,5)P2 was increased in the vesicles (**Fig. 1j, k**). Importantly, control GUVs containing no PI(4,5)P2 but only 20% PS in PC did not afford any emission in the blue channel demonstrating the distinct selectivity of **MFR-12aa** toward PI(4,5)P2 (**Fig. 1j**). This result indicated that the probe had the potential to detect PI(4,5)P2 in the cellular milieu that would have a similar membrane composition.

## MFR-12aa images PI(4,5)P2 pools in live cells and in a multi-cellular organism

We next tested whether **MFR-12aa** could permeate cell membranes and image PI(4,5)P2 in living cells. The probe entered living cells within 5-15 min of direct incubation (**Fig 1l, Fig. S7 and Fig. 4a-d)**. Importantly, the probe could enter both cultured mammalian cells like HEK293T and primary cortical neurons (**Fig 1l, Fig. S7 and Fig. S14)**. The probe did not affect cell-viability up to a tested concentration of 10 µM in HEK293T cells (**Fig. S8**) and because of its apt sensitivity could be employed at as low as 1 µM incubation levels for imaging experiments in living cells. When HEK293T cells incubated with the **MFR-12aa** probe were imaged under a fluorescence confocal microscopy setup, fluorescence emission was observed from the cytoplasmic regions of the cell, in the orange channel (580-620 nm) which represents emission from both the free and PI(4,5)P2 bound probe (**Fig. S7**). This data showed that the probe was cell-permeable. The blue channel emission (480- 520 nm) which represents emission from predominantly the PI(4,5)P2 bound probe was observed from the plasma membrane (**Fig. S7 and Fig. 1l**) implying that the probe could image plasma membrane PI(4,5)P2 pools. We hence proceeded to explore the ability of the probe to image PI(4,5)P2 pools in vivo, in a multicellular organism. We chose *C.elegans* as a model organism since these are optically transparent microscopic organisms and have a well-defined nervous system. Transgenic *C.elegans* (*gqIs25*) that overexpress PI(4)P-5-kinase (PI5K) in neuronal cells were specifically used for the experiments. The enzyme PI5K converts PI(4)P to PI(4,5)P2 and previous quantification of PI(4,5)P2 levels in the neurons of *gqIs25* showed ∼ 40% higher PI(4,5)P2 levels when compared to the neurons of wild-type *C.elegans* (*N2*) strains.^27^ Hence, this multicellular model system provided a unique handle to not only test the in vivo applicability but also importantly validate the PI(4,5)P2 selectivity of the **MFR-12aa** probe.

Live *C.elegans* were directly incubated with the probe in aqueous buffer and anesthetized prior to confocal imaging. Confocal images of the live worms showed bright fluorescence in orange (**Fig. S9**) and blue emission channels (**Fig. 2c, d**), from the mouth along with emission from the pharynx, and regions surrounding these organs for both the wild-type *N2* strain and the transgenic *gqIs25* strain (**Fig. S9**, **Fig. 2c, d**). This observation suggested that the probe primarily entered the animals through their mouth and then diffused to tissues that were in the proximity. The blue emission channel corresponding to the PI(4,5)P2-bound probe channel showed a pattern around the pharynx for the *gqIs25* strain (**Fig. 2d**) which could be identified as the ‘nerve ring’ which is a bundle of neurons that form a ring-like assembly around the pharynx (**Fig. 2a, b, d**). 3D reconstructions were used to generate a 360° view image that reconfirmed that the blue emission was significantly brighter in the nerve ring (**Fig. 2d, Video-3**) and distinguishable from the background (**Fig. 2c**). Emission from the nerve ring was observed in all the *gqIs25* worms that were imaged (*n*=18). For wild-type worms (*N2* strains), that have endogenous PI(4,5)P2 levels in neurons, the emission intensity from the nerve ring was lower (**Video-4**) than the intensity obtained for the *gqIs25* worms as expected and was observed in 75% of the worms that were imaged (**Fig. 2c,d**). Intensity analysis of the images indicated that the absolute blue channel emission intensities from the nerve ring were 40% lower in the *N2* strain compared to the *gqIs25* strain (**Fig. 2e**). The average ratio of the blue channel emission intensity from the nerve ring and background was also significantly higher for the transgenic strain by 28% when compared to the wild-type strain (**Fig. 2f**). These results distinctly proved that **MFR-12aa** was both a cell and tissue-permeable, PI(4,5)P2 selective and sensitive probe that could image PI(4,5)P2 in living cells and importantly in vivo in a multicellular organism. With this validation we next proceeded to visualize PI(4,5)P2 dynamics in the context of GPCR signaling.

**Fig 2.**
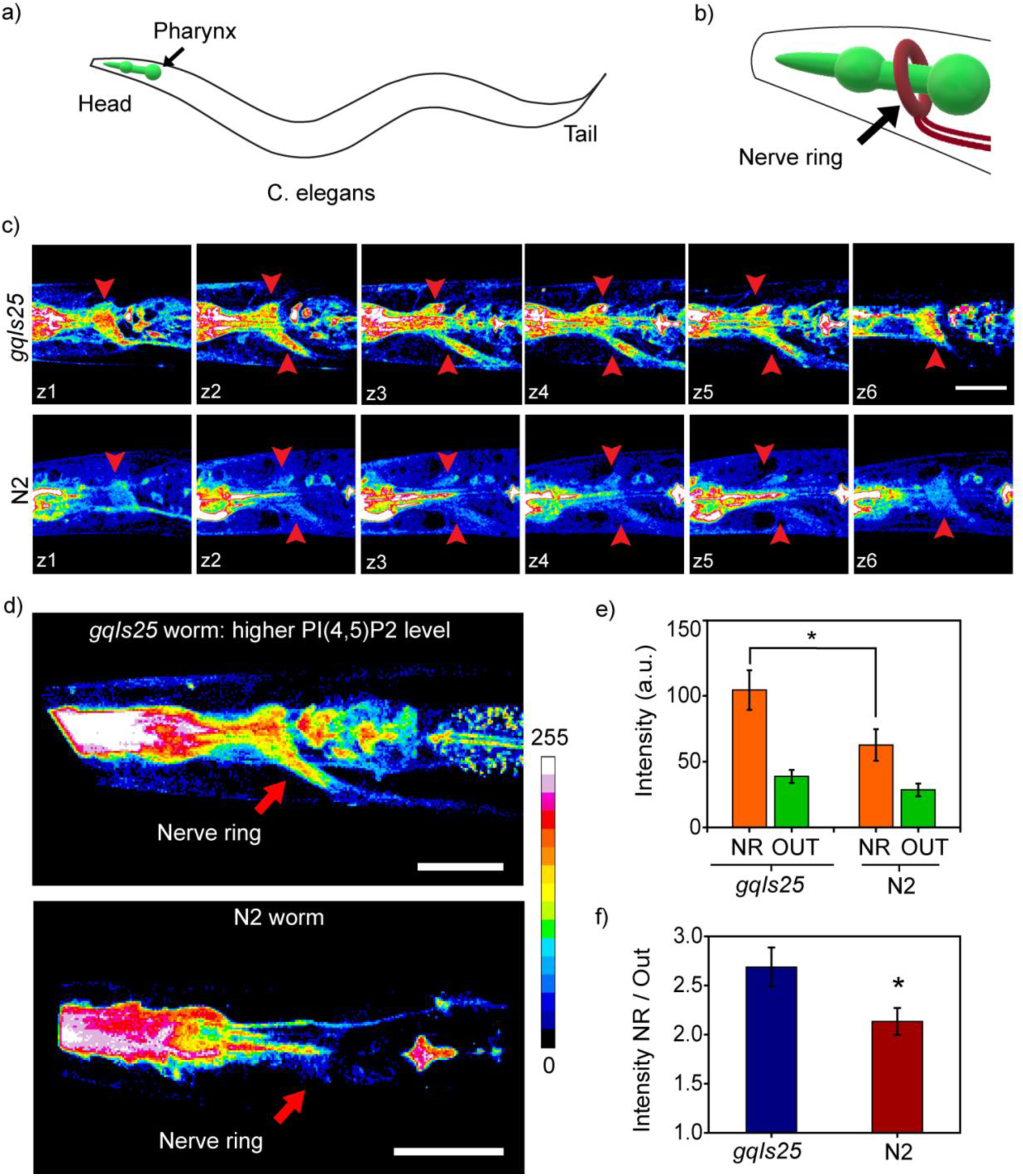
MFR-12aa images PI(4,5)P2 in a multicellular model system. (a) Diagram of *C. elegans* with pharynx region highlighted in green. (b) Enlarged diagram showing the nerve ring around the pharynx in the head region. (c) Confocal z-sections (*z1-z6*) of the head region of a PI5K overexpressing transgenic (*gqIs25*) worm (top panel) and a wild type *N2* worm (bottom panel). Images were acquired in the blue channel (480-520 nm) and the regions identified as the sections of the nerve ring are pointed with red arrows. (d) 3D reconstructions from the confocal images recorded at the head region of a PI5K overexpressing transgenic (*gqIs25*) worm (top) and wild type worm (bottom). (e) Bar plot representing raw fluorescence intensities from a region of interest (ROI) selected within the area identified as nerve ring (NR) and an ROI of same area chosen within the tissue section which is away from the pharynx and the nerve ring (OUT) for transgenic and wild type animals. (f) Ratio of intensities in the NR with respect to that of ROI outside NR. Error bars represent SEM. Unpaired Student’s *t* test, *n*=18 for transgenic and *n*=14 for *N2*, from 3 independent experiments*, *p=0.04, **p=0.03*. Scale bar, 10 µm.

## Timing PI(4,5)P2 dynamics during GPCR signaling in a cell line

With the PI(4,5)P2 sensitive and selective, small-size, cell-permeable probe we next asked if the marked difference in physiological responses to hallucinogenic versus non-hallucinogenic ligands of the 5-HT_2A_ receptor could be traced back to one of the earliest molecular events in the signaling response to these ligands, i.e. hydrolysis of PI(4,5)P2 when an agonist binds to a Gq-coupled GPCR. We focused on the 5-HT_2A_ receptor, which upon binding its natural ligand, serotonin (5 hydroxytryptamine, 5-HT), activates PLC mediated hydrolysis of PI(4,5)P2 into IP3 and diacylglycerol (DAG) (**Fig. S10)**. The 5-HT_2A_ receptor is also one of the primary targets for serotonergic psychedelics, and is thought to contribute to the diverse behavioral and physiological effects of these potent compounds. In this regard, the 5-HT_2A_ receptor provides a platform to assess the earliest lipid signaling signatures that can arise in response to the endogenous ligand, serotonin; a representative potent serotonergic psychedelic, the hallucinogenic agonist 2,5-dimethoxy-4- iodoamphetamine (DOI); or a representative non-hallucinogenic agonist Lisuride.

For the first set of experiments, HEK293 cells over-expressing 5-HT_2A_ receptor-eGFP were used. ^28^ The experimental workflow involved incubating living cells with the **MFR-12aa** probe, setting up the cells on a fluorescence confocal microscope stage and collecting time lapse images at every 5 s. The ligands were added after 60 s of collection of basal fluorescence emission, followed by ligand removal at 210 s to monitor the re- synthesis of PI(4,5)P2 at the membrane. The details of the experimental parameters are provided in the supporting information, section 15.

Addition of 5-HT to living cells, resulted in 25% depletion in the fluorescence signal of the PI(4,5)P2 bound probe blue emission channel compared to the initial time point (**Fig. 3b,c, Video-5)**. The extent of depletion observed with 5-HT should include the effect of almost simultaneous replenishment of PI(4,5)P2 on the membrane via kinase activity, which can be significantly fast during receptor-mediated signaling processes.^3^ In order to validate this point, we added a kinase inhibitor, wortmannin (WMN, 10 µM), along with 5-HT. WMN, when applied at ∼ 10 µM levels can inhibit the re-synthesis of PI(4,5)P2 from PI(4)P in the plasma membrane.^29,30^ Addition of 5-HT+WMN to living cells, resulted in 50% depletion in the fluorescence signal of the PI(4,5)P2 bound probe blue emission channel compared to the initial time point (**Fig. 3a, b, d, Video-6**). A control experiment with only WMN showed no depletion in the fluorescence intensity indicating that the depletion was specific to the loss of PI(4,5)P2 from the cell membrane upon the binding of 5-HT to the 5-HT_2A_ receptor (**Video-5**). Notably, upon ligand wash for the 5-HT+WMN treated cells, we observed a compensation of the depleted signal from the plasma membrane (**Fig. 3a, b, d, Video-6**) which demonstrated the reversibility of the probe response and its distinct ability in tracking PI(4,5)P2 dynamics in living cells. The signal intensity, however, did not recover back to the original level which could be due to very little loss of unbound probe, if any, during multiple washing steps. This was also reflected in the 5-HT treated cells in which the signal did not recover to the initial starting intensity upon wash. No significant loss in fluorescence intensity across the plasma membrane was observed in the case of vehicle-treated cells, when monitored for the same time duration precluding any photo-damage of the probe and establishing that **MFR-12aa** was photo-stable (**Fig. 3b, Video-6**). Next, we proceeded to visualize PI(4,5)P2 dynamics in response to a hallucinogenic versus a non-hallucinogenic agonist of the 5-HT_2A_ receptor.

**Fig 3.**
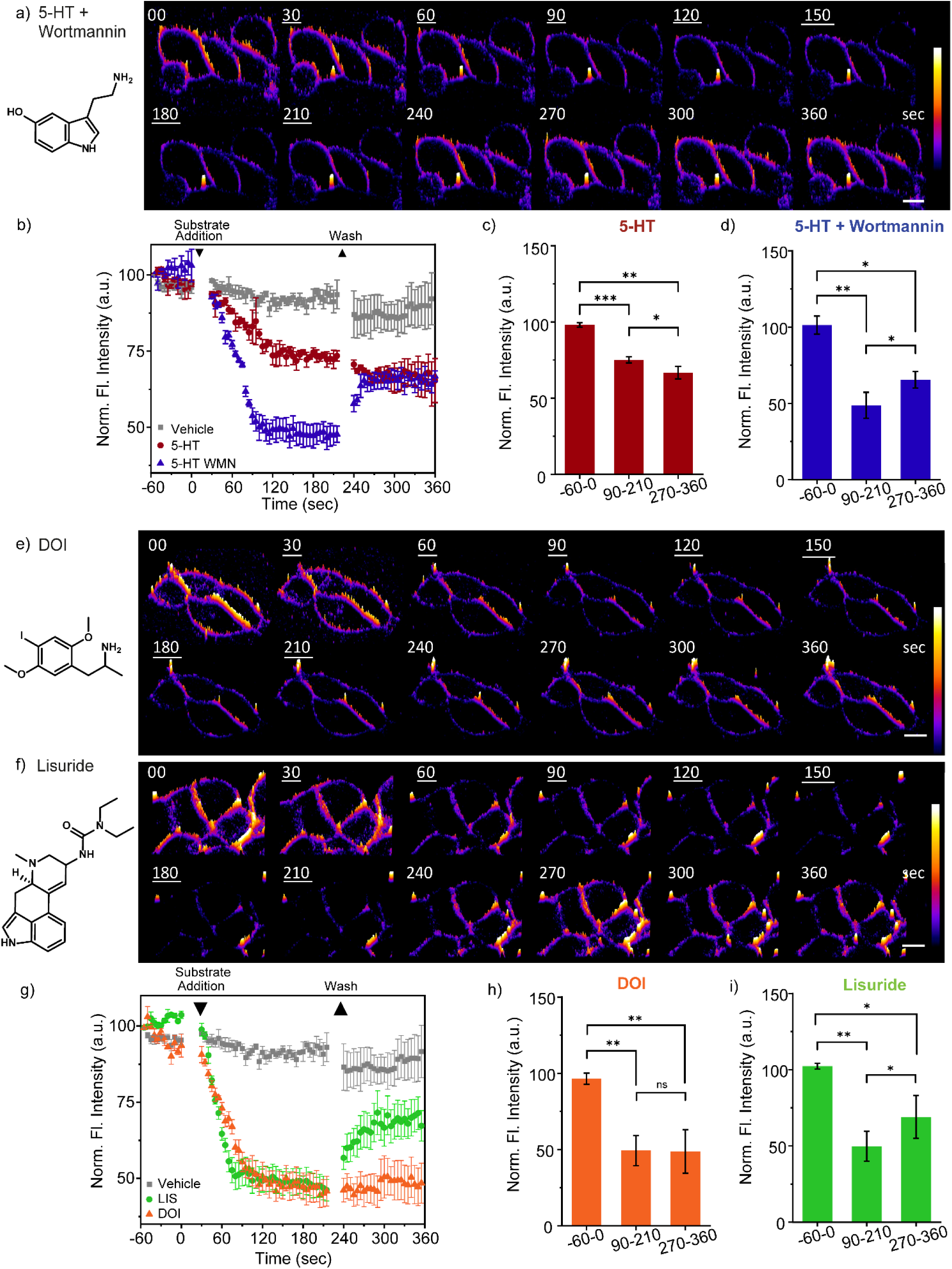
Neurotransmitter, hallucinogen, and non-hallucinogen mediated PI(4,5)P2 dynamics in live 5-HT_2A_ receptor over-expressing HEK293 cells. Confocal time lapse imaging of cells incubated with **MFR-12aa** (5 µM) using two-photon excitation λ_ex_ 780 nm, λ_em_ 480-520 nm. Scale bar, 10 μm. Time trace for a single z-slice in 3D intensity plot representation during stimulation of cells with: (a) 5-HT (10 μM) and WMN (10 μM); (e) DOI (10 μM); and (f) Lisuride (10 μM). Data were acquired for unstimulated cells (1 min, -60-0 s), followed by stimulation via ligand addition at 0 s (3 min, 30-210 s) and ligand removal via replacement with fresh media at 210 s (3 min, 240-360 s). Time points indicating the start and end of stimulation are underlined. Data depicted is representative of three independent repeats for each experiment. Time traces of fluorescence intensity, calculated by normalizing the fluorescence intensity, depicting differential rates of ligand dependent PI(4,5)P2 hydrolysis and replenishment for: (b) 5-HT (maroon), 5-HT + WMN (blue), and vehicle (0.01% DMSO) (gray), and (g) DOI (orange), Lisuride (green), and vehicle (0.01% DMSO) (gray). Fluorescence intensity in the blue channel in the selected ROI of the membrane from individual experiment was averaged and normalized to the initial intensity (t= -60 s) of the same experiment. Mean of these averaged values from three independent experiments are plotted as a time trace until 360 s of image acquisition. Error bars represent standard error of mean from three individual experiments. Normalized fluorescence intensity bar plots representing average intensity values for the time intervals -60-0 s, 90-210 s, 270-360 s calculated from time lapse images of cells treated with: (c) 5-HT (10 μM); (d) 5-HT (10 μM) and WMN (10 μM); (h) DOI (10 μM); and (i) Lisuride (10 μM). Ligand addition has been marked at 0 s and washed at 210 s. Error bars represent standard error of mean from three individual experiments (*N* = 3; vehicle, *n* = 46; 5-HT, *n* = 46; 5-HT and WMN, *n* = 47; DOI, *n* = 41; LIS, *n* = 47); *\* p = 0.02, ** p = 0.01, *** p = 0.001* using Student’s *t-*test.

DOI the hallucinogenic agonist, has a higher affinity to the 5-HT_2A_ receptor, with respect to the endogenous ligand 5-HT.^31^ Upon addition of DOI to living cells followed by time-lapse imaging, a ∼ 50% depletion of the blue channel emission of the probe was observed (**Fig. 3e, g, h**). The PI(4,5)P2 depletion response was significantly larger than that observed in the case of 5-HT stimulation alone and comparable to the stimulation by 5-HT with WMN (**Fig. 3b, g**). For, DOI and 5-HT+WMN, the signal from the blue channel decreased steadily till 95 s post ligand addition and then plateaued. On the other hand, stimulation with Lisuride, a non-hallucinogenic agonist, that binds the receptor with comparable affinity as DOI,^32^ could bring about faster PI(4,5)P2 depletion within 75 s, reaching a similar plateau as that of DOI or 5-HT+WMN (**Fig. 3g**, **Fig. S12a**). Surprisingly, we observed no replenishment of PI(4,5)P2 upon washing the agonist in case of DOI-treated cells (**Fig. 3e, g, h, Video-7**) while the Lisuride treated cells afforded recovery of the blue emission intensity of the probe upon agonist washing, indicating PI(4,5)P2 replenishment within the timescale of the imaging experiment **(Fig. 3f, g, i, Video-7**). The result was robust and reproducible and indicated significant ligand-specific differences in PI(4,5)P2 temporal dynamics within 50-100 s post agonist addition (**Fig. S12a**).

To test whether these early signaling event differences propagated to downstream processes, we independently monitored two downstream events which follow the action of PLC on PI(4,5)P2: 1. Receptor, 5- HT_2A_ receptor-eGFP, endocytosis which occurs almost immediately following ligand-binding;^28^ 2. Amounts of inositol-monophosphate (IP1), a stable metabolite of IP3. Additionally, the levels of diacylglycerol (DAG), the product of PI(4,5)P2 depletion on the cell membrane were also measured.

When 5-HT was added to the cells, 5-HT_2A_ receptors internalized within a time frame of 110-115 s (**Fig. S11, Video-8**), validating our probe response data depicting PI(4,5)P2 hydrolysis saturation within a similar time range (105-115 s) (**Fig. 3b**). In the case of 5-HT+WMN, the extent of receptor internalization was similar, depicting an unbiased response of 5-HT towards 5-HT_2A_ receptor internalization, even in the presence of WMN (**Fig. S11e, f**). In a key observation, the difference in PI(4,5)P2 depletion rates were also reflected in the rates of receptor endocytosis, with Lisuride showing faster receptor internalization than DOI (**Fig. S11c-f, Video-9, Fig. S12**). Replenishment of the GFP-tagged 5-HT_2A_ receptor back to the plasma membrane at the end of the stimulation, has been reported to be a rather slow process (120-180 min)^33^ and thus mirroring the replenishment, in this case, was not performed.

Next, IP1 production in DOI and Lisuride stimulated HEK293-5-HT_2A_ receptor-eGFP cells, was examined using ELISA (**Fig. S13**). End-stage IP1 and DAG levels for Lisuride stimulated cells remain lower in magnitude across the doses and time points tested, as compared to DOI. These results plausibly indicate conversion of IP1 and DAG into replenishing PI(4,5)P2 pools back to the cellular membranes, in line with earlier reported observations.^34^ Overall our results indicate that the **MFR-12aa** probe can time differences in one of the earliest signaling events post-ligand binding to a GPCR and that these early time differences are robust and propagate through downstream signaling paths. We next investigated whether the differences observed in PI(4,5)P2 dynamics in the 5-HT_2A_ receptor overexpression cell model could be observed in primary neurons which are the actual site of action for these signaling processes.

## Tracking temporal PI(4,5)P2 dynamics in primary cortical neurons using MFR-12aa

Successful demonstration of the **MFR-12aa** probe toward timing and tracking PI(4,5)P2 dynamics in a 5-HT_2A_ receptor expressing stable cell line, encouraged us to test the probe in the context of primary cortical neurons expressing endogenous levels of 5-HT_2A_ receptor in the cell body and neurites.^35^ Primary neurons upon incubation with 1 µM of **MFR-12aa** showed blue/bound probe emission from the membrane of the cell body as well as the neurites (**Fig. 4a-d, Fig. S14**), within 5 min of direct incubation.^36^ This result indicated the distinct ability of the probe to directly permeate not only cultured cells but also primary neurons, which is a rare feature in molecular probes.

Upon stimulation with 5-HT alone, we observed an ∼8 % greater depletion in the PI(4,5)P2 bound probe blue emission from the neurites (26 % depletion) in comparison to the cell body (18 % depletion ) (**Fig. 4b, e-h; Video-10**). The differential depletion might be attributed to the higher expression of the 5-HT_2A_ receptor in the neurites, and indicate the possibility that the probe could detect for local cellular differences in active receptor signaling pools in neurons.^37^ The extents of signal depletion were comparable to 5-HT mediated stimulation alone in the HEK293-5-HT_2A_ over-expression system (25% depletion) (**Fig. 3b, c**). In the case of DOI, we observed a 52% depletion of the probe response from the neurites and 31% depletion from the cell body that were significantly higher as compared to 5-HT stimulation (**Fig. 4c, e-h, Video-11**). On the other hand, in the case of Lisuride-treated primary neurons (**Fig. 4d, e-h, Video-11**), the depletion of signals from the neurites was 42% and cell body was 22% which was lower than that of DOI stimulated neurons, but higher than that evoked by 5-HT stimulation. To illustrate the differential temporal effects of hallucinogenic versus non-hallucinogenic agonists, DOI and Lisuride, snapshots from the early time points have been shown in **Fig. S15** where we note a significant reduction in the blue-channel emission intensity of the sensor within 30 s of DOI addition, whereas in the case of Lisuride the extent of intensity decrease at the same time-point is lower (**Video-11**). Together the data distinctly provide evidence for the significantly longer-lasting effects of DOI, the potent hallucinogenic agonist, in comparison to Lisuride, the non-hallucinogenic agonist, at the very earliest stages of signaling events, like PI(4,5)P2 depletion.

**Fig 4.**
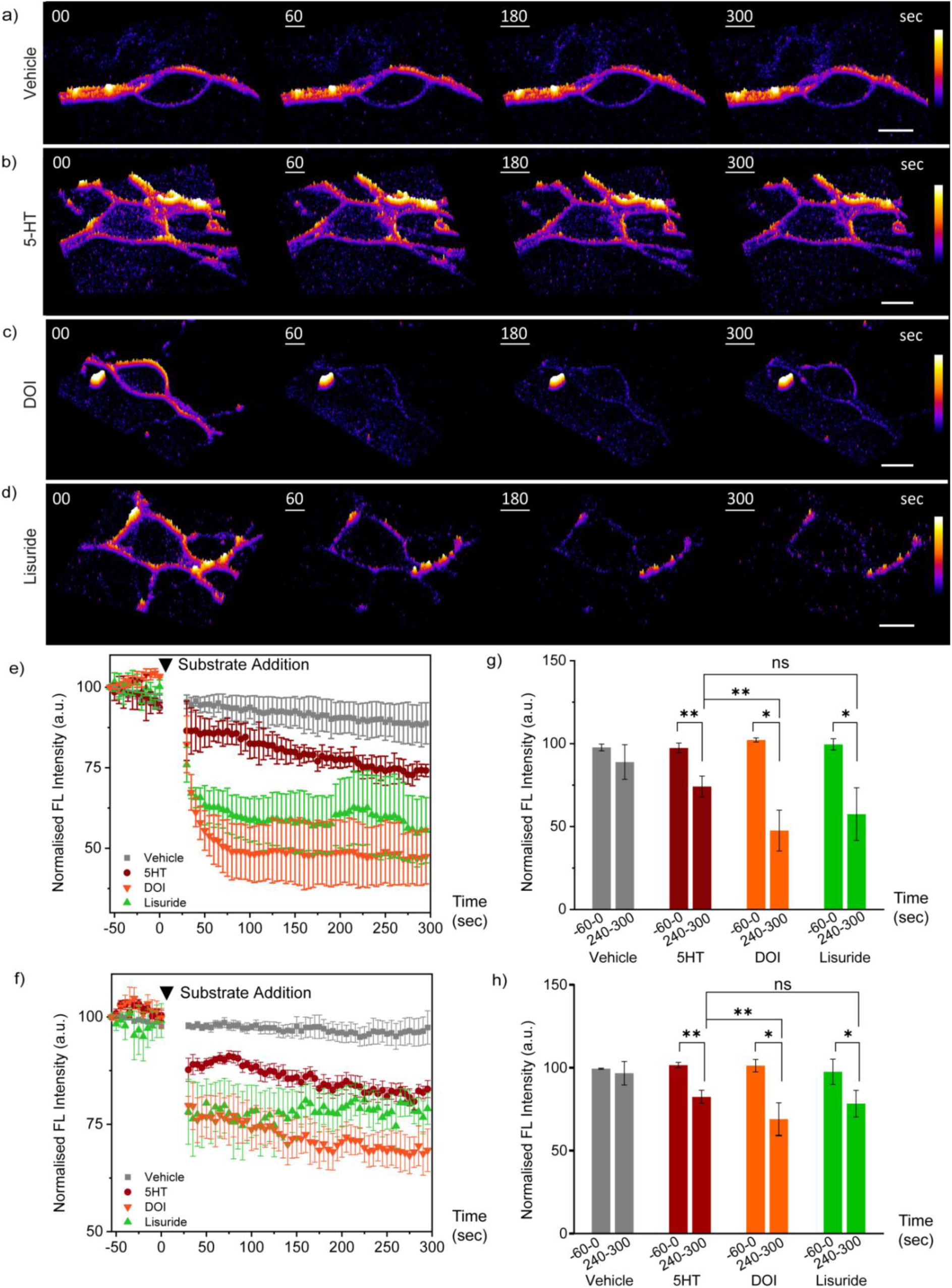
Localized dynamics in cortical neurons in presence of neurotransmitter, hallucinogen, and non-hallucinogenic agonists. (a-d) Confocal time lapse imaging of rat primary cortical neurons incubated with **MFR-12aa** (1 µM), using two-photon excitation at λ_ex_ 780 nm, λ_em_ 480-520 nm. Time trace for a single z-slice in 3D intensity plot representation during stimulation of cells with: (a) vehicle (0.01% DMSO); (b) 5-HT (10 µM); (c) DOI (10 µM); and (d) Lisuride (10 µM). Data were acquired for unstimulated cells (1 min, t= -60-0 s), followed by stimulation via ligand addition at 0 s (5 min, 0-300 s). Time points indicating stimulation are underlined. *N*=4. (e,f) Time traces of fluorescence intensity depicting differential rates of ligand dependent PI(4,5)P2 hydrolysis for: 5-HT, DOI, Lisuride, and vehicle (0.01% DMSO), for: (e) neurites; and (f) neuron cell body. Fluorescence intensity in the blue channel in the selected ROI on the membrane from individual experiment was averaged and normalized to the initial intensity (t=-60 s) of the same experiment. Mean of these averaged values from three independent experiments are plotted as a time trace until 300 s of image acquisition. Time points indicating ligand addition are pointed by downward arrowheads. Normalized fluorescence intensity bar plots representing average intensity values for the time intervals -60-0 s, and 240-360 s calculated from time lapse images of cells treated with 5-HT (10 µM), DOI (10 µM), Lisuride (10 µM), and vehicle for (g) neurites and (h) neuron cell body. Error bars represent SEM, *N=4* (cell body: *n* = 3-5: cell processes: *n* = 11-15); *\* p* = 0.02, *\*\* p* = 0.01, *\*\*\*p*=0.001 using Student’s t-test, scale bar, 10 μm.

These findings provide evidence for distinct signaling signatures evoked by hallucinogenic versus non- hallucinogenic 5-HT_2A_ receptor agonists at the earliest stages of signal initiation, uncovered through visualization of the differential rates of PI(4,5)P2 hydrolysis revealed by our probe. These unique results provide evidence of the heterogeneity in signaling to distinct ligands at the same receptor, not only in terms of the nature of time course, but also the manner of signal activation. Our results from cell lines and primary neurons, show that our cell-permeable reversible probe can discern subtle PI(4,5)P2 dynamic signatures in living systems.

## Perspective

It is well-established that different ligands of the same Gq-coupled GPCR evoke diverse physiological responses; yet, a key unanswered question is if these signaling outcomes are linked to early-time point differences at the signal initiation stage. To address this question, we need to time early molecular steps that occur within seconds of ligand binding. However, tracking seconds-minute temporal differences in signal transduction, live, has been a significant challenge owing to the lack of apt chemical tools. There is a critical need to reversibly and non-invasively track specific phosphoinositides which are amongst the first molecules at the cell membrane that respond to and mediate signals following the binding of a ligand to a Gq-coupled GPCR. We have addressed this challenge by developing a peptide-based, cell-permeable, ratiometric fluorescent probe **MFR-12aa**. In vitro, in cell, and in vivo studies in a multicellular organism validated that the probe could selectively and sensitively detect PI(4,5)P2, the phosphoinositide that mediates Gq-coupled GPCR signaling. The probe has two key salient features: First, reversibility, which was introduced into the probe design by strategically converting non-covalent interactions between the peptide-binding scaffold and PI(4,5)P2 into a fluorescence response; Second, photo-stability, achieved via a fluorene-based fluorophore. These features played a pivotal role as we interrogated whether the responses of a hallucinogenic versus a non-hallucinogenic agonist of the serotonergic receptor showed any differences in PI(4,5)P2 mediated signaling within seconds of agonist-receptor binding. In a central result of this report, we show that the probe could indeed time differences in PI(4,5)P2 depletion, an early time point signaling event that occurs upon ligand binding to a Gq-coupled GPCR, and capture differential temporal dynamics of PI(4,5)P2 for a hallucinogenic versus a non-hallucinogenic agonist of the 5-HT_2A_ receptor. Given the highly differential behavioral and physiological outcomes when the 5-HT_2A_ receptor is stimulated by hallucinogenic versus non- hallucinogenic agonists, our results provide evidence of distinctive changes in PI(4,5)P2 depletion, that are revealed at the early stages of ligand-receptor interaction. This result opens the floodgates for applying short- peptide based cell-permeable lipid probes for tracking the spatiotemporal dynamics of these critical signal mediators. The power of this tool lies in uncovering the earliest of signaling signatures that may underlie the distinctive physiological and behavioral outcomes that arise in responses to diverse ligands of Gq-coupled receptors.

## Supporting information

video-description

## Acknowledgments

The authors thank the cell culture facility, confocal facility, and MALDI facility at the Department of Chemical Sciences, TIFR. We acknowledge the kind gift of the human 5-HT2A receptor-EGFP fusion receptor expressing HEK293 cell lines from Dr. Mitradas Panicker, National Centre for Biological Sciences, Bangalore, India. A.D. acknowledges support from the Department of Atomic Energy, Government of India, under Project Identification No. RTI4003. A.S.K acknowledges support from Agence Nationale de la Recherche (ANR) AmpliSens ANR-21-CE42-0019-01.

## Author Contributions

AD guided, designed, conceptualized, and oversaw the project; RK and SM performed the simulations, designed and developed probes, conducted in vitro experiments, and analyzed the data; AK and AAB conducted experiments in cells and primary cortical neurons and analyzed data; SPK assisted in planning and setting up *C.elegans* experiments and SM performed the experiments; VAV assisted in setting up and planning neuronal experiments; RV assisted in molecular dynamics simulations and analysis; OAK and ASK synthesized and shared polarity-sensitive dye; RK, SM, AK, AAB, VAV, and AD wrote the paper. All authors reviewed the results and the manuscript and approved the final version of the manuscript.

