## Supplementary Information for "A Cell-Permeable Fluorescent Probe Reveals Temporally Diverse PI(4,5)P2 Dynamics Evoked by Distinct GPCR Agonists in Neurons"

### Supplementary methods.

#### 1. Simulation system preparation

Simulations were performed using the NAMD 2.11 simulation package. Initial peptide coordinates were obtained from RCSB protein data bank, PDB ID 1SOL. The lipid bilayer model was generated by VMD software using the 'Membrane Builder' plug-in. A membrane of 100 Å x 100 Å dimension composed of POPC was built using CHARMM36 force field. The coordinate files for PI(4)P (SAPI14), PI(4,5)P2 (SAPI24) and PI(3,4,5)P3 (SAPI33) were taken from the "CHARMM-GUI Archive - Individual Lipid Molecule Library". In each case, a single PI(4)P/ PI(4,5)P2/ PI(3,4,5)P3 molecule was inserted into one leaflet of the POPC membrane. For each phosphoinositide we generated a model comprising of one phosphoinositide molecule per 135 POPC molecules which corresponded to 0.74 mol% of phosphoinositide on that leaflet. The missing atoms were added using CHARMM36 force field for lipids as well as peptide structure and psf files were generated from the pdb file inputs. The pdb and psf files were used for equilibration, minimization, and production runs. After generation of the lipid bilayers, the model systems were solvated in a water box followed by neutralization and ionization with $Na^+$ and $Cl^-$ ions maintaining the NaCl concentration at 120 mM. The equilibrated peptide was incorporated into the system after the lipid bilayer was equilibrated. All the simulations were run with a periodic simulation box condition.

#### 2. Simulation details

*Equilibration of the membrane.* Membrane equilibrations were performed using the NAMD simulation package and CHARMM36 force field. The lipid bilayer was energy minimized and equilibrated through 3 steps – a) 10000 steps of minimization followed by 0.5 ns equilibration at 300 K keeping everything free except the lipid tails fixed; b) Repetition of minimization and equilibration steps keeping all non-hydrogen atoms under harmonic constrains (changing from higher to lower) both lipid tails and headgroups, water and ions free. c) The whole system was equilibrated again at 300 K for 1 ns setting all atoms free.

*Equilibration of the peptide.* The peptide structure was decided on the basis of previously reported literature which proposed that, upon binding to PI(4,5)P2 the Gel-20aa sequence formed a helix, imposing a major conformation change in the protein, displacing it from actin filaments.[1] The structure of the 20aa peptide was obtained from Protein Data Bank (PDB ID: 1SOL) which was acquired via an NMR experiment in presence of helix inducing solvent TFE.[2] The peptide PDB structure had a $–COO^-$ group at the C-terminal. As the experiments were performed with $–CONH_2$ terminal peptides, the C-terminal capping was modified to $–CONH_2$

by using a patch in VMD before equilibration and minimization. The peptide was incorporated into a modeled water box generated via VMD. The box size was maintained such that the distance between the peptide and its periodic image remained more than 28 Å in all directions. The solvent box was neutralized by adding the requisite number of $Na^+$ and $Cl^-$ ions and total NaCl concentration was maintained to be 120 mM. The system was subjected to the following minimization steps: (a) keeping the non-hydrogen heavy atoms in the peptide fixed with harmonic constraints of 25 and 12 kcal/mol/$Å^2$ on the peptide heavy atoms; (b) The minimization steps were followed by heating the system from 0 K to 300 K in two steps: (1) steady increment of temperature in steps of 6 K/1 ps from 0 K to 300 K; (2) equilibration at 300 K for 50 ps keeping the protein heavy atoms fixed; (3) repeating the heating and equilibrium steps keeping the heavy atoms under harmonic constraints of 25 kcal/mol/$Å^2$. The harmonic constraints were gradually released in NPT (1 atm pressure and 300 K temperature) equilibration steps of 1 ns. For the shorter peptide, Gel-12aa, structure was generated by truncating the equilibrated 20aa peptide structure after the 12$^{th}$ amino acid from the C-terminus. Then the N-terminus was capped with $-CONH_2$. Rest of the steps were done following the above-mentioned procedures.

*Insertion of the peptide onto the lipid bilayer and simulation.* The equilibrated peptide was aligned with the equilibrated membrane containing PI(4,5)P2 in two different orientations: a) keeping the C-terminus close to the headgroup, and b) keeping the N-terminus close to the headgroup. In both cases upon running short 100 ns simulations, stable electrostatic contacts were formed between the C-terminus positively charged residues and the phosphoinositide headgroup. Hence, for all following simulations, the C-terminus was placed close to the phosphoinositide headgroup at the starting point of the simulations with the membrane. The peptide-membrane system was re-solvated and ionized with 120 mM NaCl. The whole system was subjected to the following steps– (a) The harmonic constraints were gradually released in five NPT (1 atm pressure and 300 K temperature) equilibration steps of 1 ns each, (b) NPT production runs were carried out for 200 ns. All simulations employed periodic boundary conditions with the PME algorithm describing long range electrostatic interactions. The van der Waals forces were calculated by using a switching function that had 10 Å as the switching distance and 12 Å as the cutoff.

Analysis of simulations was conducted using NAMD. N-terminal dipping, non-bonding energy, and peptide helicity calculations were done on the .dcd files generated after the final NPT production run. For 200 ns trajectories, we have considered 1000 frames for all the calculations. The distance (z-coordinate difference) between the N-terminus amine group of the peptide and the membrane surface, calculated throughout the simulation, determined the peptide's position relative to the membrane, with positive values indicating above-surface placement and negative values indicating penetration into the membrane. The non-bonding energies were computed between positively charged amino acid residues on the peptide, which approached within 3 Å of the phosphoinositide headgroup and negatively charged phosphate groups on the phosphoinositide headgroup for the calculations.

A summary of simulations performed is provided in **table S2**.

**3. Peptide synthesis**

Solid-phase peptide synthesis resins, amino acids, and activating reagents used in the peptide synthesis were procured from Novabiochem (Merck Millipore). Peptides were synthesized by using solid phase peptide synthesis strategy on Fmoc protected Rink amide resin HL (100-200 mesh, loading 0.74 mmol/g resin) using an automated peptide synthesizer (PS3, Protein technologies Inc., USA). Rink amide resin (100 mg) was de-protected using 30% (v/v) piperidine in dimethylformamide (DMF). For each coupling step, Fmoc protected amino acid (3 eq., 0.22 mmol), either 3-[Bis(dimethylamino)methyliumyl]-3H-benzotriazol-1-oxide hexafluorophosphate (HBTU) or 1-[Bis(dimethylamino)-methylene]-1H-1,2,3-triazolo-[4,5b]-pyridinium-3-oxide- hexafluorophosphate (HATU), (3 eq., 0.22 mmol), and N-methylmorpholine (4% (v/v) in DMF) were used. After the final Fmoc deprotection, the peptides conjugated to the beads were washed with DMF and MeOH and stored under $N_2$ atmosphere at -20 ˚C. In order to cleave the peptides from the beads, the beads were treated with a cleavage cocktail having trifluoroacetic acid, triisopropyl-silane, phenol, water, ethanedithiol, and thioanisole (in 80: 2.5: 5: 5: 2.5: 5 volume ratio) for 4 h. The resins were removed from the mixture and the solution was concentrated under a stream of $N_2$. The crude peptides were precipitated out in cold methyl tert-butyl ether. The peptides were then characterized by LC-ESI-MS (LCMS-2020, Shimadzu Corp) and MALDI-TOF-MS (Bruker UltrafleXtreme, Bruker Daltonics).

**4. Synthesis of MFR-dye.**

***7-Bromo-9,9-dimethylfluorenyl-2-amine*** was obtained as described previously.[3]

***7-bromo-N,N,9,9-tetramethyl-9H-fluoren-2-amine***

To a stirred mixture of 7-bromo-9,9-dimethylfluorenyl-2-amine (750 mg, 2.60 mmol) and paraformaldehyde (780 mg, 26.0 mmol) in 99 % glacial acetic acid (10 mL) at 25°C under nitrogen atmosphere, $NaCNBH_3$ (820 mg, 13.0 mmol) was added in one portion. The resulting mixture was stirred at room temperature for 24 h, and then it was poured into cold water (150 mL). The resultant solution was extracted with ethyl acetate (2×30 mL). The combined organic layer was washed with saturated aq. $NaHCO_3$ solution and dried with sodium sulfate. Organic solvent was removed under reduced pressure, and the obtained residue was purified by column chromatography (eluent $CH_2Cl_2$:Heptane = 1:1) to give the desired product as a white powder. Yield was 620 mg (75 %). Product could be crystallized from ethyl alcohol.

$^1$H NMR ($CDCl_3$, 400 MHz): δ 7.55 (d, 1H), 7.49 (d, 1H), 7.43 (d, 1H), 7.40 (dd, 1H), 6.76 (d, 1H), 6.72 (dd, 1H), 3.04 (s, 6H), 1.46 (s, 6H).

$^{13}$C NMR ($CDCl_3$, 100 MHz): δ 27.36, 40.87, 46.95, 106.32, 111.54, 118.62, 119.86, 120.76, 125.71, 129.79, 138.87, 154.91, 154.93.

LCMS: (m/z): MM-ES+APCI, found $[M+1]^+$ 316.1 (calculated for $C_{17}H_{18}BrN+H^+$ =316.1)

***4-(7-(dimethylamino)-9,9-dimethyl-9H-fluoren-2-yl)-4-oxobutanoic acid (MFR-COOH)***

7-bromo-N,N,9,9-tetramethyl-9H-fluoren-2-amine (620 mg, 1.96 mmol) and magnesium turnings (52 mg, 2.16 mmol) were transformed into Grignard reagent upon heating at 50 °C in dry THF for 1 h. The obtained clear solution was cooled and added dropwise to the solution of succinic anhydride (392 mg, 3.92 mmol) in THF at -78 °C under nitrogen atmosphere. After the addition was completed, the reaction mixture was allowed to heat to room temperature, and then it was stirred for additional 4 h. The reaction was quenched with 10 % aq. HCl solution, diluted with 30 mL of distilled water and extracted with chloroform (2×30 mL). The combined organic layer was dried with $Na_2SO_4$ and evaporated under vacuum. The obtained residue was purified by column chromatography (eluent $CH_2Cl_2$:MeOH = 99:1, then 98:2, the 2$^{nd}$ fraction was collected) to afford the final compound as a yellow powder. Yield was 200 mg (30 %).

$^1$H NMR (400 MHz, $CDCl_3$ with 10 % Methanol-$d_4$): δ 7.99 (d, 1H), 7.93 (dd, 1H), 7.62 (d, 1H), 7.59 (d, 1H), 6.75 (d, 1H), 6.73 (dd, 1H), 3.36 (t, 2H), 3.05 (s, 6H), 2.82 (t, 2H), 1.47 (s, 6H).

$^{13}$C NMR (100 MHz, $CDCl_3$+10 % Methanol-$d_4$): δ 27.28, 28.25, 33.15, 40.76, 46.82, 106.10, 111.64, 118.08, 121.91, 121.92, 126.55, 128.10, 133.24, 145.37, 150.85, 151.40, 152.98, 156.85, 197.51.

HRMS (m/z): ESI, found $[M+1]^+$ = 338.175 (calculated for $C_{21}H_{23}NO_3+H^+$ = 338.175).

**5. Labelling of the peptide with MFR-COOH dye**

Labelling with MFR-COOH dye was performed on the peptides conjugated to beads. Synthesized peptide on beads (20 mg) was coupled with MFR-COOH (3 eq., 0.22 mmol), HATU (3 eq., 0.22 mmol), and N, N-Diisopropylethylamine (DIPEA) (3 eq., 0.22 mmol) by mechanical mixing for 48 h. Labelled peptide on the beads was washed with DMF to remove the excess unreacted dyes and beads were dried under vacuum. Peptide was cleaved from the resin by treating with the cleavage cocktail mentioned previously, for 4 h. The resin beads were removed from the mixture and the solution was concentrated under a stream of $N_2$. The crude peptides were precipitated out in cold methyl tert-butyl ether.

**6. Purification and characterization of the MFR probes**

MFR labeled peptides were purified on a Varian PrepStar, SD-1 HPLC system, equipped with a Phenomenex Luna® C18(2) (250 x 10 mm) semi-preparative column. Water (solvent A) with trifluoroaceticacid (0.1% v/v) and acetonitrile (solvent B) with trifluoroaceticacid (0.1% v/v) were used as mobile phase and absorption at 220 nm and 390 nm were monitored for product elution. The purification method was set to a gradient flow for 0-5 min: 5% solvent B; 5-60 min: 5-100% solvent B; 60-65 min: 100% solvent B; 65-75 min: 100-5% solvent B; 80 min: 5% solvent B. HPLC fractions were characterized by MALDI-TOF-MS (Bruker UltrafleXtreme, Bruker Daltonics) using HCCA matrix to identify fractions containing the desired product. Pure fractions were evaporated under reduced pressure on a vacuum concentrator (Eppendorf Concentrator Plus, Eppendorf AG, Germany). Purity of the probes were further confirmed by electro-spray ionization liquid chromatography mass spectroscopy (LC-ESI/MS) and MALDI-TOF-MS.

**7. SUV preparation.**

All the lipids were procured from Avanti Polar Lipids, Inc. Phospholipids were taken in a glass vial in the desired molar ratio and mixed either in chloroform or in a mixture of chloroform: methanol: water (20:9:1). After mixing by vortex, the solvents were evaporated first under $N_2$ and then overnight under vacuum to obtain lipid films. Lipid films were then hydrated in buffer (20 mM Na-HEPES, 100 mM NaCl, pH 7.4) and taken through 5 freeze-thaw cycles using liquid nitrogen and a hot water bath. SUVs were prepared via extrusion by passing the aqueous suspensions through polycarbonate membranes of 100 nm pore diameter at 25 °C. Formation of vesicles was confirmed by size measurements using Dynamic Light Scattering (DynaPro, S9 Protein Solutions, Wyatt Technology Corp., USA). Concentrations of the phospholipid stock solutions were determined by estimating the total phosphate content using previously reported procedure.[4,5]

**8. Fluorescence titration of the probes with small unilamellar vesicles (SUVs)**

All measurements were performed in buffer (20 mM HEPES, 100 mM NaCl, pH 7.4) at 25 °C. Probes (1 µM) were dissolved in buffer and concentrations of the stock solutions were measured using a NanoDrop 1000 spectrophotometer (Thermo Fisher Scientific, Inc.) by monitoring the MFR absorbance at 390 nm (molar absorptivity of MFR at 390 nm = 43000 $M^{-1}cm^{-1}$). Probe solutions (1 µM, 200 µL) were mixed with the SUVs and fluorescence spectra were recorded in a FluoroLog-3 spectrofluorometer (Horiba Jobin Yvon, Inc.) by excitation at 390 nm. To determine dissociation constants ($K_d$), probes were titrated against SUVs composed of different phospholipids and $F_{480}/F_{580}$ values were plotted against corresponding phospholipid concentrations. The response curves were fitted to the equation, $(F_{480}/F_{580}) = (F_{480}/F_{580})_{max}/(1+K_d/[phospholipid])+(F_{480}/F_{580})_0$ for ratiometric fitting for **MFR-20aa** and $F= (F_{max}-F)_0/(1+K_d/[phospholipid])+F_0$ for turn on fitting for **MFR-12aa** and $K_d$ values were obtained from the fits. For comparison studies, fluorescence spectra were recorded with the probes (1 µM) in the presence of either SUVs or IP3.

**9. GUV preparation.**

GUVs were prepared according to a previously reported method.[6] In brief, lipid stock solutions containing phospholipids in desired molar ratio (keeping total phospholipid concentration 1 mg/mL) were prepared in chloroform. Clean glass coverslips were placed on a hot plate pre-heated to 75 °C and aqueous poly(vinyl alcohol) (PVA) solution (5% w/v, 25 µL) was spread on each coverslip. The coverslips were left for drying. This procedure led to the formation of a thin film of PVA on the coverslips. Lipid stock solutions (3 µL) were placed on the PVA films. After the solvent evaporated completely, the films were peeled off and immersed in 150 µL of buffer (20 mM Na-HEPES, 100 mM NaCl, pH 7.4) and left undisturbed for 30-40 minutes at room temperature. The GUVs formed were released from the PVA film by aspiration using a pipette and collected in fresh microtubes.

**10. Confocal imaging of probes on giant unilamellar vesicles (GUVs)**

GUVs were incubated in 200 nM probe solution and allowed to settle for 10 min. Imaging was done on a Ziess 710 multiphoton confocal microscope. MaiTai Titanium Sapphire Crystal laser was set at 780 nm and power adjusted optimally. GUV imaging was done on a 40x water immersion objective with N/A of ¼. A suitable position with spatially oriented GUV was selected and imaged. $\lambda_{ex}$ was set at 780 nm and images were recorded. Blue (bound) 480-520 nm channel images have been shown in Fig. 1j. Laser power and detector gain were adjusted in order to avoid signal saturation at the highest PI(4,5)P2 concentration imaged. All GUV images were recorded under identical laser power and detector gain. Confocal images were analyzed using ImageJ (NIH, USA) software and the intensities along the perimeter of the confocal images of the GUVs were measured using a Concentric Circles plugin. Three independent experiments were performed and 10 GUVs were considered for analysis in each experiment.

**11. Imaging PI(4,5)P2 in HEK293T cells**

HEK293T cells were cultured in Dulbecco's Modified Eagle Medium (DMEM, Sigma-Aldrich) high glucose supplemented with fetal bovine serum (10%), Penicillin (50 units/mL) and Streptomycin (50 µg/mL) in T-25 culture plates at 37 °C inside an incubator containing humidified 5% $CO_2$. For imaging, the cells were seeded on home-made glass coverslip bottomed dishes (35 mm diameter, Tarsons) coated with polylysine (0.1 mg/mL), and incubated at 37 °C under 5% $CO_2$ for 24 h. Cells plated on glass coverslips were washed with DMEM high glucose, phenol red free medium and incubated at 37 °C with **MFR-12aa** (5 µM in phenol red free DMEM) for 15 min. Following incubation, the cells were washed twice with phenol red free DMEM. $\lambda_{ex}$ was set at 780 nm and images were recorded in two different channels, blue (bound) 480-520 nm and orange (bound+unbound) 580-620 nm.

**12. Imaging phosphoinositides in *C. elegans***

Wild type (N2) and gqIs25 strains of *C. elegans* were grown following standard protocols. All imaging experiments were performed at the young adult stages (between L4 to 1-day adult) of the worms. Before probe incubation worms were washed with Milli-Q water. The washed worms were then transferred into a solution containing **MFR-12aa** (10 µM) in M9 buffer and were allowed to swim freely for 1 h at room temperature. After the incubation, worms were taken out of the probe solution and again washed with water to remove excess probe from the body surface. For confocal imaging, worms were anesthetized with sodium azide solution (30 mM, in M9 buffer, 10 min). The anesthetized worms were placed onto an agar pad (2% low density agar in Milli-Q water) made on a microscope slide and gently covered with a glass cover-slip. The slide containing worms was placed on the microscope stage keeping the cover-slip side facing the microscope objective. Worms were imaged with a 40x water immersion objective upon two-photon excitation at 780 nm using the same instrument settings that were used for cellular imaging with **MFR-12aa**. Optical sections along the z-direction were taken at every 1 µM difference in order to create 3D reconstructions. Confocal images were processed and analyzed using ImageJ software.

**13. Imaging phosphoinositide dynamics in HEK293 cells over-expressing 5-HT$_{2A}$ receptor-eGFP**

Generation and characterization of HEK293 cells stably expressing the human serotonin receptor type 2A fused to eGFP (HEK293- 5-HT$_{2A}$ receptor-eGFP cells) has been described previously.[41] Cells were maintained in

DMEM high glucose supplemented with 10% FBS, antibiotic-antimycotic solution and neomycin analogue G418 (1 mg/ml) in T25 culture flasks at 37°C, 5% $CO_2$. 24 hours prior to imaging studies, cells were seeded (2 x $10^5$ cells per well) in 35-mm glass bottom dishes (Nunc). Cells were initially starved with DMEM without FBS supplemented for 4-6 hours incubated at 37 °C. Serum free media was replaced with **MFR-12aa** probe solution (5 µM) in DMEM high glucose, phenol red free medium for 15 min incubated at 37 °C. Probe solution was washed twice with Phenol red free DMEM and imaged on a Ziess 710 Multiphoton Confocal Microscope.

**14. Primary cortical neuronal cultures**

Primary cortical neuronal cultures were established from postnatal day 0 Sprague Dawley rat pups. All experimental procedures were in accordance with the guidelines of the Committee for Supervision and Care of Experimental Animals (CPCSEA), Government of India, and were approved by the TIFR Institutional Animal Ethics committee.

Imaging dishes were pre-coated 4-6 hours prior to seeding with 0.1 mg/mL Poly-D-Lysine hydrobromide. Under aseptic conditions, pup brains were dissected and placed in Hanks' balanced salt solution supplemented with antibiotic-antimycotic solution. The cortical tissue was dissected and trypsinized for 10 min at 37°C. Thereafter, cells were dissociated by repeated trituration using Minimum Essential Medium Eagle (MEM) containing Hank's Salts, 25 mM HEPES buffer, L-Glutamine and Sodium bicarbonate supplemented with 10% FBS. Cells were then plated (2x$10^5$ cells per well) in DMEM supplemented with 10% FBS at 37°C, 5% $CO_2$ for 3-4 h. After that, the media was replaced to Neurobasal medium, supplemented with 2% B27, 0.5 mM L-Glutamine and antibiotic-antimycotic solution. Half of the medium in the dishes was replaced every alternative day with fresh neurobasal medium containing the supplements mentioned above. Cells were maintained for 7-10 days *in vitro* before use in experiments and experiments were performed when the cultures comprised of 90-95% neurons. For vehicle or drug inductions prior to imaging, Neurobasal phenol red free medium was used. During probe incubation, supplemented Neurobasal media was replaced with **MFR-12aa** probe solution (1 µM) in Neurobasal, phenol red free medium for 5 mins and incubated at 37 °C. Probe solution was washed twice with phenol red free Neurobasal and imaged on a Ziess 710 Multiphoton Confocal Microscope.

**15. Confocal Imaging parameters**

Imaging was performed on a Ziess 710 Multiphoton Confocal Microscope. MaiTai Titanium Sapphire Crystal laser was set at 780 nm and adjusted to a power of ~ 30 mW on a 10x objective. Cell Imaging was done on a 40x water immersion objective with N/A of ¼. A suitable position with spatially and uniformly distributed cells was chosen on the plate and the best focused z-slice was set. $\lambda_{ex}$ was set at 780 nm and images wwere procured in two different channels, blue (bound) 480-520 nm and orange (bound+unbound) 580-620 nm using an external PMT for the transmission image. Three independent experiments were performed and a total of 30-50 cells were considered for analysis. A timeline was followed for capturing the depletion and replenishment response of the probe in a stipulated time as mentioned below. Data acquisition was started in a cycle mode of 5 s interval for a total time of 6 min. An initial 60 s (12 frames) was acquired to record the intracellular/endogenous flux of PI(4,5)P2 dynamics. Acquisition was paused and media was replaced with the respective concentration of the desired ligands. Removal and addition of ligands took around 20-22 s; thus a time delay of 25 s was maintained using a stopwatch, wherein the data acquisition was resumed again. After 180 s of ligand exposure, data acquisition was paused again, (210 s) ligand solution was removed, and fresh media was added. This was repeated twice to ensure complete removal of ligand. Again, data acquisition was resumed after 30 s (240 s), for an additional time of 2 min. For primary neuronal cells data acquisition was continued until 300 s of ligand exposure.

**16. Estimation of DOI/ Lisuride induced intracellular secondary messengers (IP1 and DAG) production by ELISA.**

HEK293-5-$HT_{2A}$ receptor-eGFP cells were seeded ($4*10^5$ cells/well) in a 24-well plate. 24 h post seeding, cells were either stimulated with the vehicle 0.1% DMSO (basal, 0) or with increasing concentrations (1 µM, 5 µM, 10 µM) of DOI/Lisuride for IP1 assay and (1 µM and 10 µM) DOI/Lisuride for DAG assay for one hour at 37°C in the incubator. After one hour, cell lysates were prepared and IP1 assay (cat. no. 72IP1PEA/D, CisBio Bioassays, Codolet, France) and DAG assay (cat. no. EKU08737, Biomatik Corporation, Ontario, Canada) were performed as per manufacturer's instructions by ELISA method. Absorbance was read at 450 nm for both the assays and the results represent mean ± SEM of three independent experiments performed in duplicates. Statistical analysis has been performed by one-way ANOVA for comparison across multiple groups followed by Tukey's post hoc test of the Software 'GraphPad Prism 5.0' (GraphPad Software, Inc., San Diego, CA, USA). The values of *$p$<0.05 with respect to basal values were considered to be statistically significant.

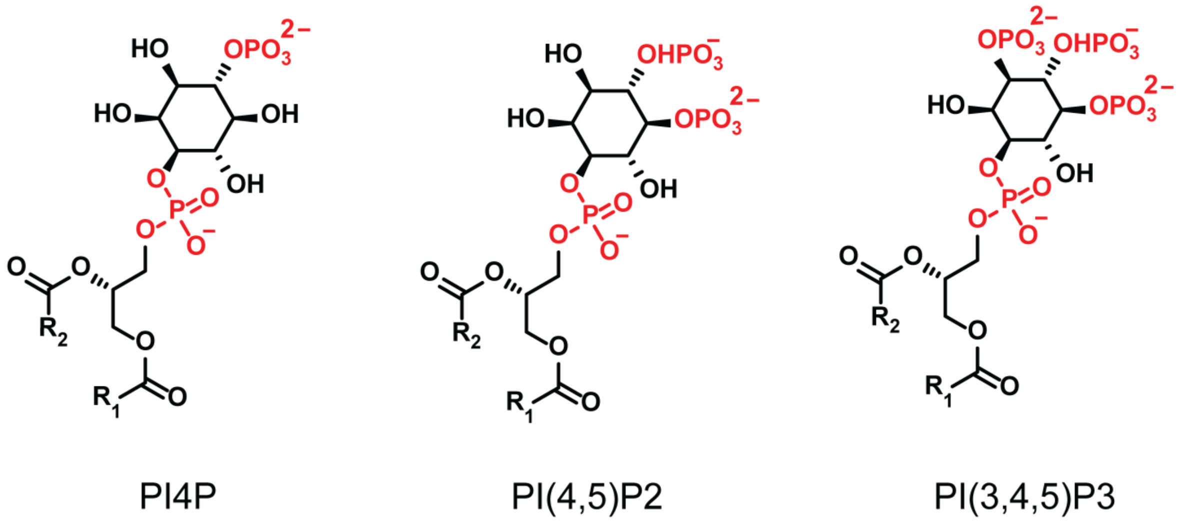


**Fig. S1|** Representative chemical structures of three major phosphoinositides out of seven variants: phosphatidylinositol-4-phosphate (PI(4)P), phosphatidylinositol-4,5-bisphosphate (PI(4,5)P2), and phosphatidylinositol-3,4,5-trisphosphate (PI(3,4,5)P3). R1 and R2 represent stearic acid and arachidonic acid tails, respectively.

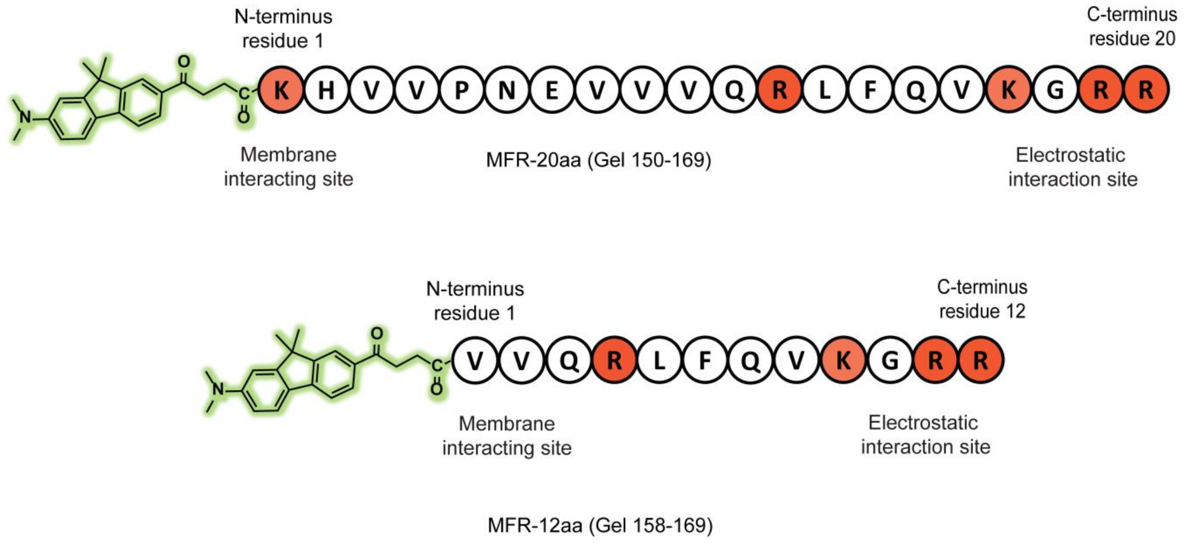


**Fig. S2|** Representative images of the probes **MFR-20aa** (above) and **MFR-12aa** (below). Cationic amino acids have been indicated in orange.

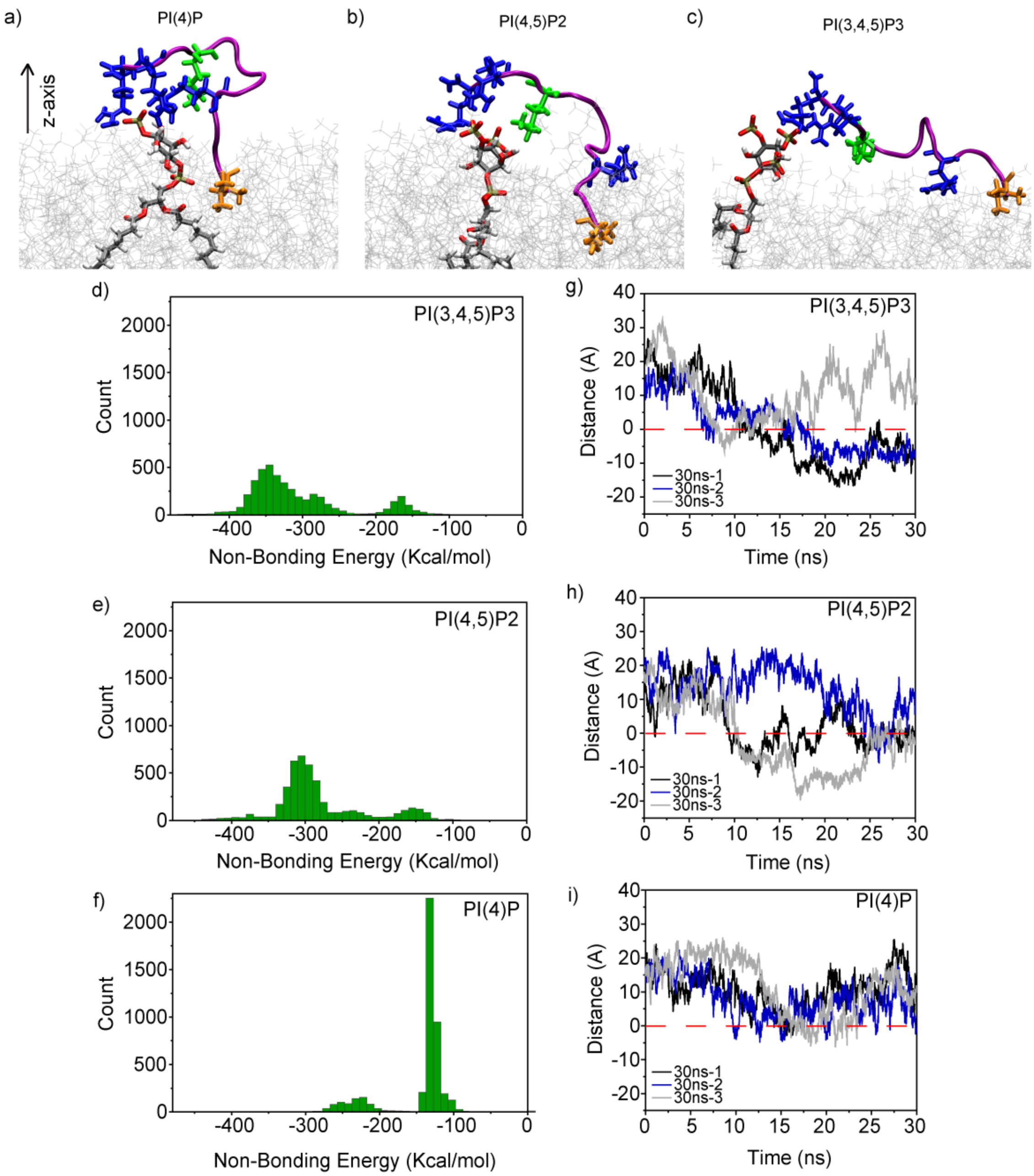


**Fig. S3|** Snapshots from MD simulations showing Gel-12aa-phosphoinositide interactions (phosphoinositide gray licorice, O-atoms in red) in a PC (gray lines) membrane bilayer: (a) PI(4)P, (b) PI(4,5)P2, and (c) PI(3,4,5)P3. The peptide backbone is in magenta and cationic amino acids are shown in blue (Arg) and green (Lys). The N-terminus amino acid has been highlighted in bronze. Distributions of non-bonding energies for binding of the Gel-12aa peptide to (d) PI(3,4,5)P3 (e) PI(4,5)P2, and (f) PI(4)P headgroups. Distributions of the distances between the N-terminus amine group of Gel-12aa and the membrane surface (dashed line) along the z-axis (**Fig. S3a**) after interacting with (g) PI(3,4,5)P3 (h) PI(4,5)P2, and (i) PI(4)P headgroups.

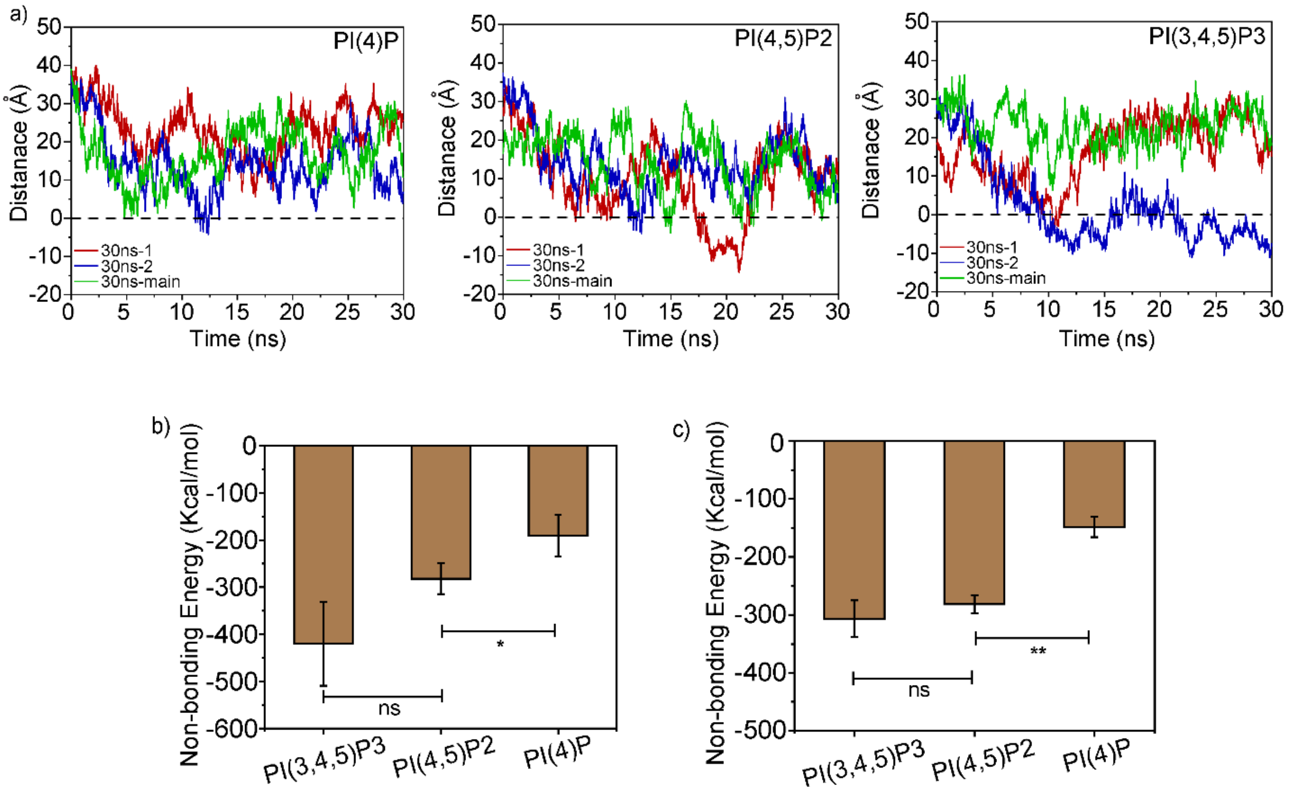


**Fig. S4|** (a) Distributions of the distance between the N-terminus amine group of Gel-20aa and the membrane surface along the z-axis upon binding to PI(4)P, (PI(4,5)P2, and PI(3,4,5)P3 headgroups, respectively. All calculations were done on two independent 30 ns simulations along with the first 30 ns taken from the 200 ns simulation for Gel-20aa-phosphoinositide-membrane systems (See **Table S2** for list of simulations). (b) Bar plot representing the average non-bonding energies for phosphoinositide headgroups and Gel-20aa, for two 30 ns simulations along with first 30 ns taken from the 200 ns simulation. (c) Bar plot representing the average non-bonding energies for phosphoinositide headgroups and Gel-12aa, for three 30 ns simulations. Error bars represent SEM. Statistical analyses were done on 3 independent sets of simulations using an unpaired, two-tailed student's t-test, $*p < 0.05$; $**p < 0.01$.

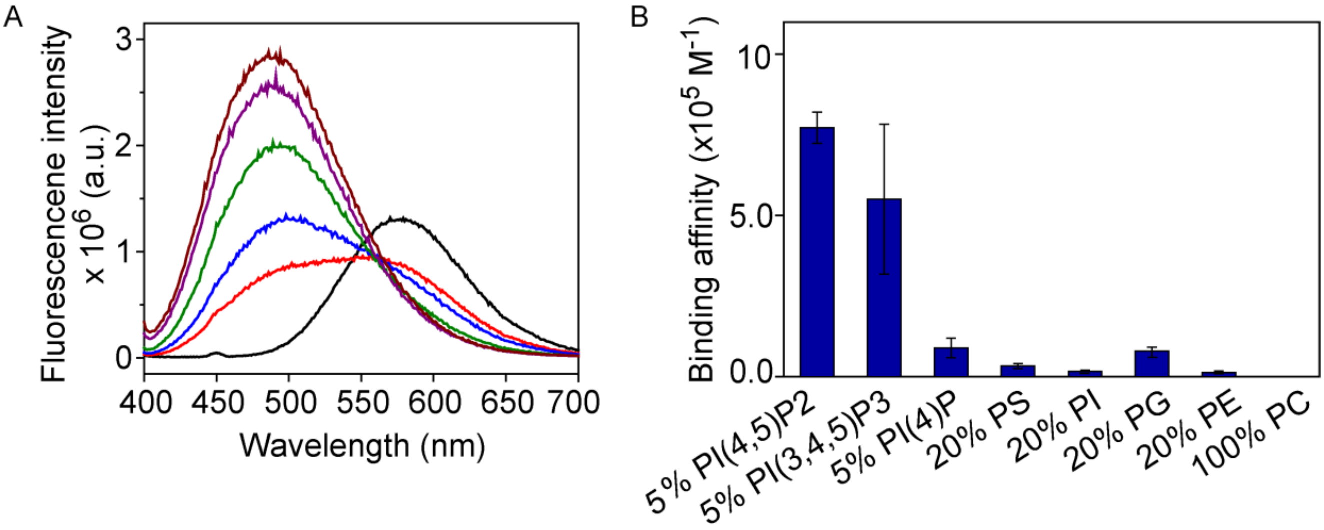


**Fig. S5|** (a) Fluorescence response of **MFR-20aa** (1 µM) depicting a blue shift in emission maxima with increasing concentrations (0, 3, 5, 24, 90, and 113 µM) of 5% PI(4,5)P2-PC SUVs, $\lambda_{ex}$ 390 nm. (b) Bar plot comparing the association constants ($K_a$) of **MFR-20aa** toward various PLs, derived from fluorescence titrations. Error bars represent SEM, $N$=3.

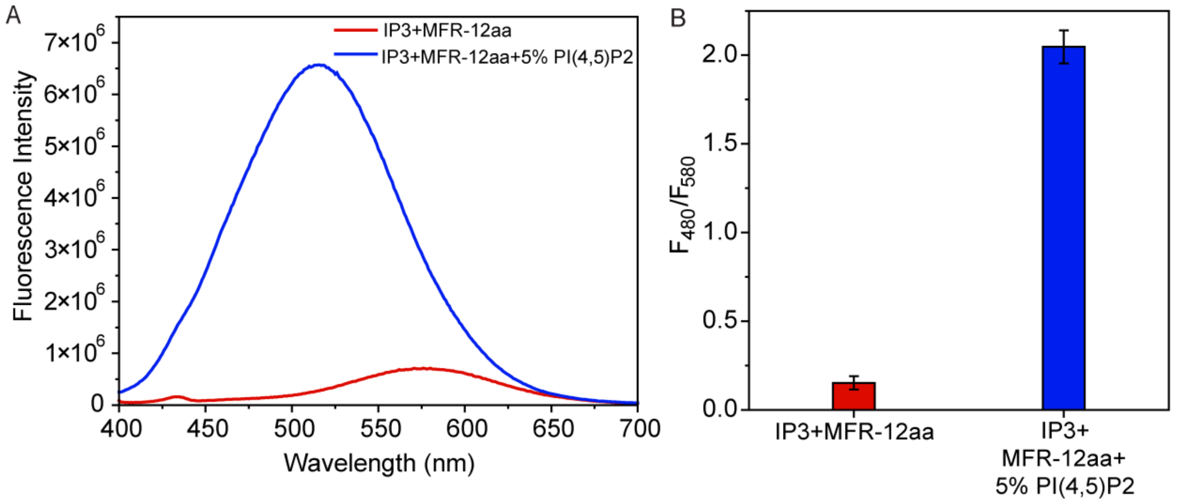


**Fig. S6|** (A) Emission spectra of **MFR-12aa** in presence of IP3 (35 µM, red) and in presence of 5% PI(4,5)P2 (70 µM total PL) +IP3 (35 µM, blue), $\lambda_{ex}$ 390 nm. (B) Ratiometric response ($F_{480}/F_{580}$) of **MFR-12aa** toward IP3 (red) and PI(4,5)P2 (70 µM total PL)+IP3 (35 µM, blue).

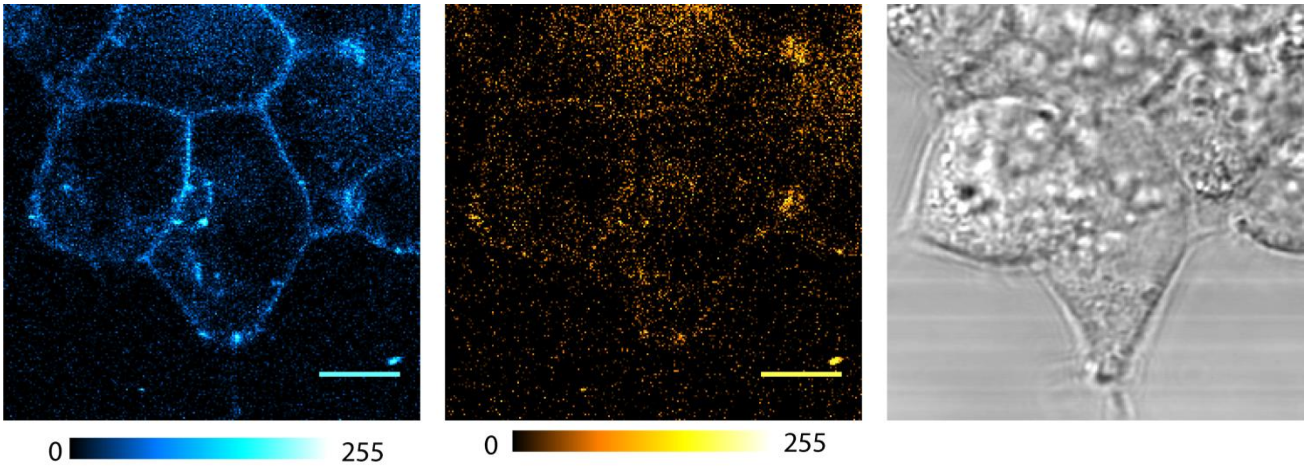


**Fig. S7|** Representative confocal single z-plane images of living HEK293T cells incubated with **MFR-12aa** (5 µM) for 15 min at room temperature. Scale bars, 10 µm. Fluorescence emission: blue channel ($\lambda_{em}$: 480−520 nm); orange channel ($\lambda_{em}$: 580−620 nm), $\lambda_{ex}$ 780 nm.

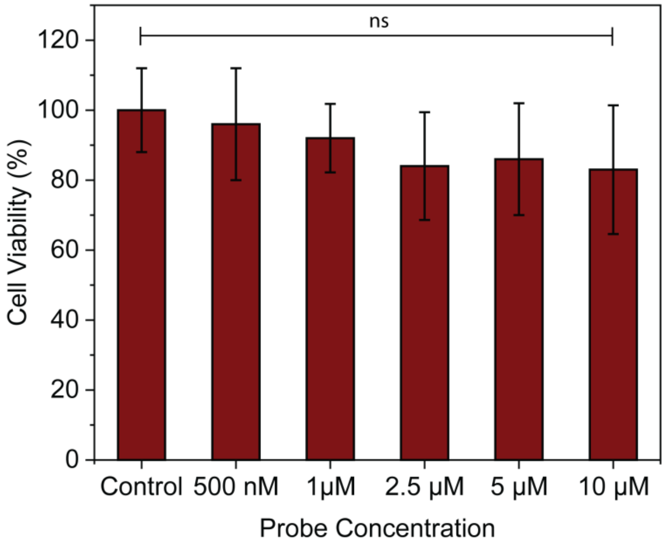


**Fig. S8|** Percentage cell viability in the presence of **MFR-12aa** at different concentrations (0, 0.5, 1, 2.5, 5 and 10 µM) in HEK293T cells after 30 min of incubation at 37 °C in DMEM with no phenol red and serum. Percentage viability was determined by recording absorbance at 570 nm for formazan using MTT assay. Data has been expressed as the mean ± SEM of three independent experiments performed in triplicates, *n*=9, *N*=3.

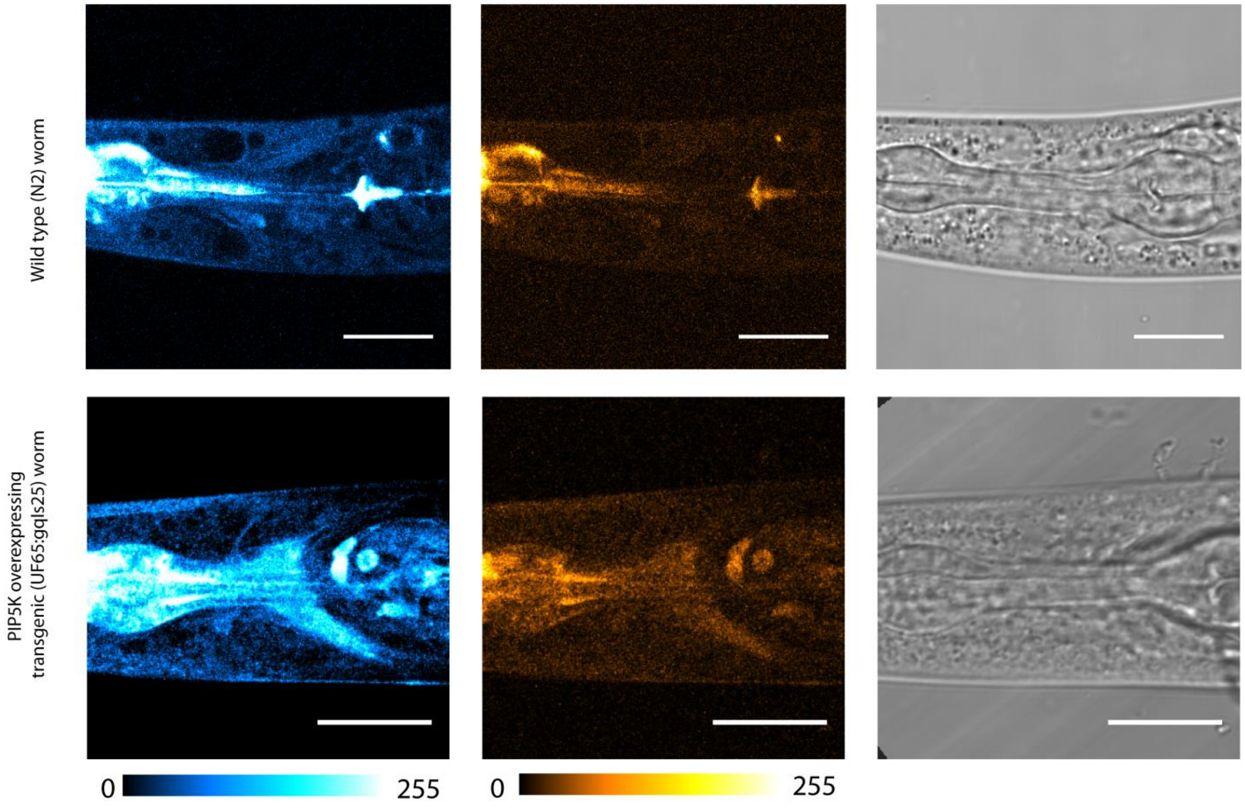


**Fig. S9|** Representative single z-plane images of live *C. elegans,* wild type *N2* (above) and transgenic *gqls25* (below). **MFR-12aa** (10 µM) was excited with 780 nm two-photon laser. Fluorescence emission: blue channel ($\lambda_{em}$: 480−520 nm); orange channel ($\lambda_{em}$: 580−620 nm).

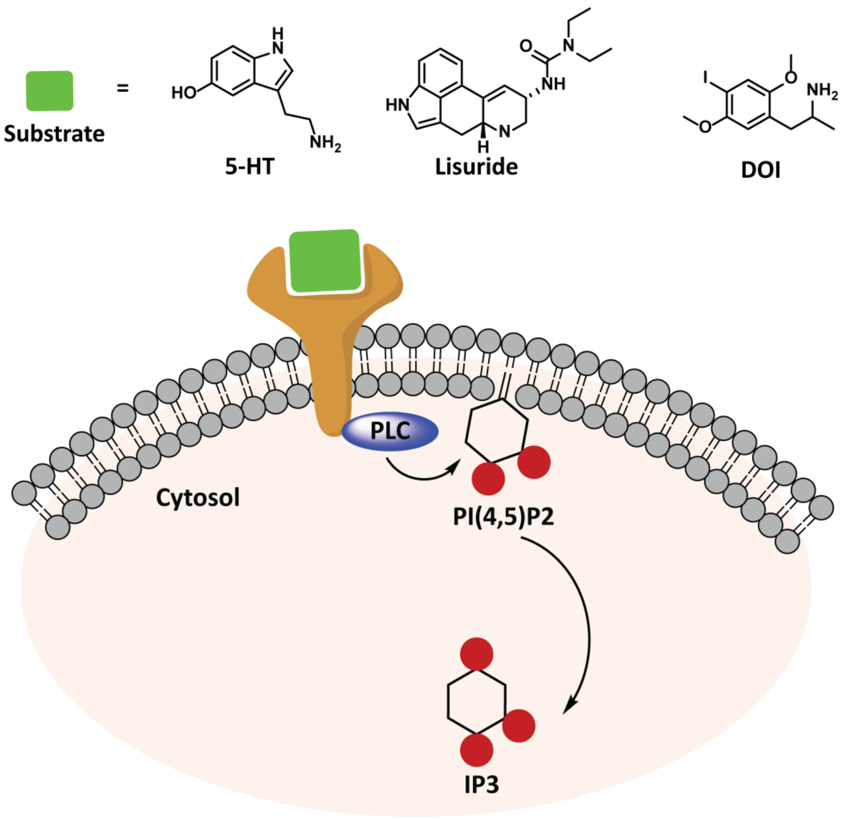


**Fig. S10|** Scheme depicting ligand mediated PI(4,5)P2 dynamics in the serotonergic receptor (5-HT$_{2A}$) system. Serotonin (5-HT), hallucinogenic (DOI), and non-hallucinogenic (Lisuride) ligands bind to the 5-HT$_{2A}$ receptor leading to activation of phospholipase C (PLC) which cleaves off the PI(4,5)P2 headgroup into IP3 (in cytosol) and DAG (in membrane). Probe binding to PI(4,5)P2 headgroup elicits a response in the membrane which diminishes upon cleavage of PI(4,5)P2 headgroup to IP3. Reversible nature of the probe aids in detection of both depletion and regeneration of PI(4,5)P2 in cellular membranes.

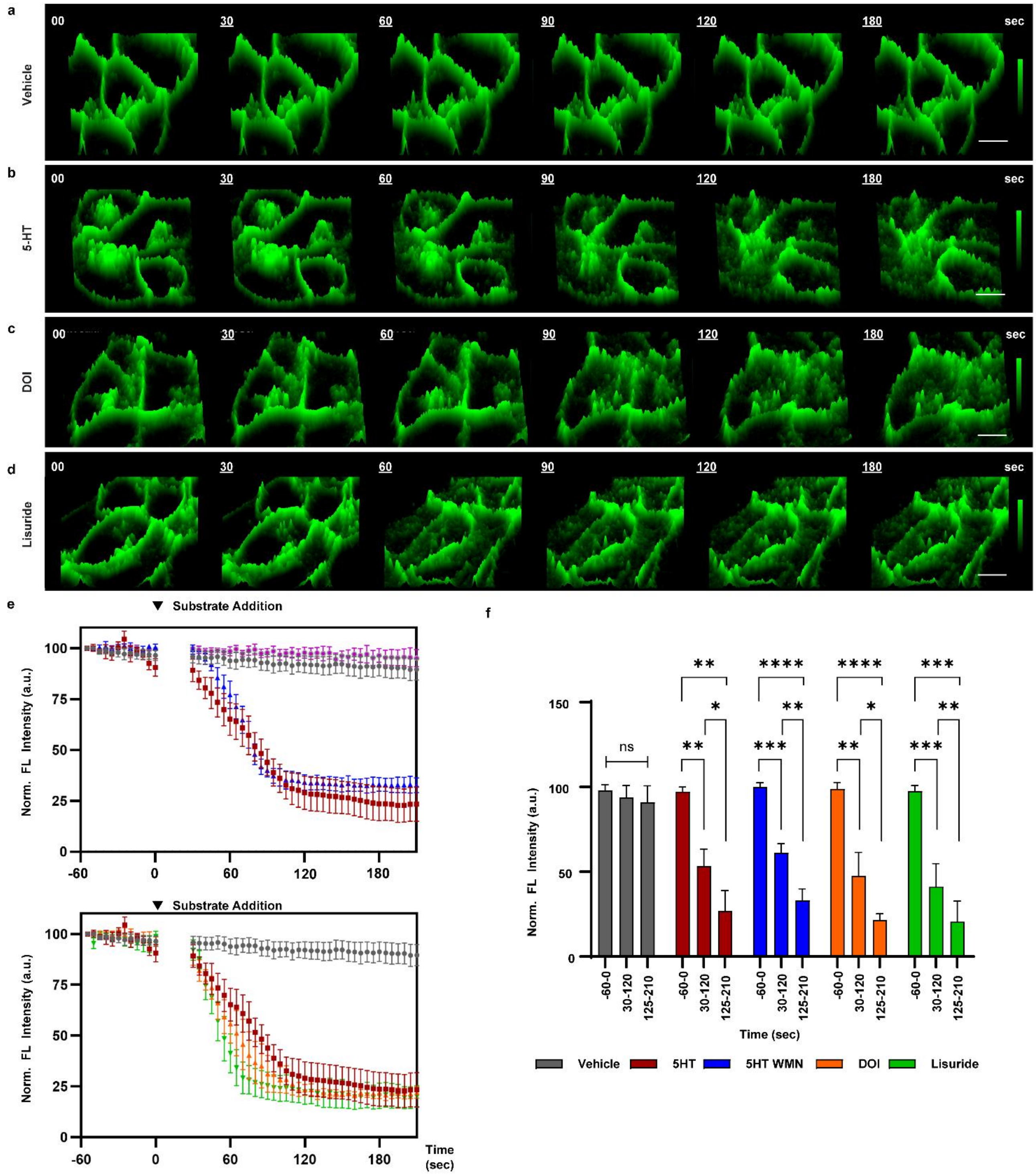


**Fig. S11**| Visualizing ligand-mediated GFP-tagged 5-$HT_{2A}$ receptor dynamics in live 5-$HT_{2A}$ over-expressing HEK293 cells using confocal time-lapse imaging in the absence of the **MFR-12aa** probe; confocal microscopy $\lambda_{ex}$ 488 nm, $\lambda_{em}$ 500-560 nm, scale bar 10 µm. Time trace for a single z-slice, 3D intensity scale representation (0-255) during (a) vehicle and ligand stimulation of (b) 5-HT (10 µM), (c) DOI (10 µM) and (d) Lisuride (10 µM). Data was acquired for unstimulated cells (1 min), ligand stimulation (3 min). Time points during ligand stimulation are underlined. Depicted data is representative of four independent experiments. (e) Top: Time trace of GFP fluorescence intensities depicting differential rates of ligand-dependent 5-$HT_{2A}$ receptor internalization upon addition of: nothing (pink); vehicle (0.01% DMSO), (gray); 5-HT (red); and 5-HT and WMN (blue). Bottom: Time trace of GFP fluorescence intensities depicting differential rates of ligand-dependent 5-$HT_{2A}$ receptor internalization upon addition of: vehicle (0.01% DMSO), (gray); 5-HT (red); DOI (Orange); and Lisuride (Green). Percentage intensities were calculated as the ratio of membrane GFP intensity to cytoplasmic GFP intensity, normalized to the initial intensity ratio. (f) Time-dependent intensity analysis bar plots showing the differential internalization rates in the absence and presence of vehicle (no internalization), 5-HT, 5-HT + WMN, DOI, and Lisuride (10 μM each, respectively). Error bars represent SEM, $N$=4 (total no. of cells = 30-50 cells); * $p$ = 0.02, ** $p$ = 0.01, *** $p$ = 0.001, **** p = 0.0001 using Student's t test.

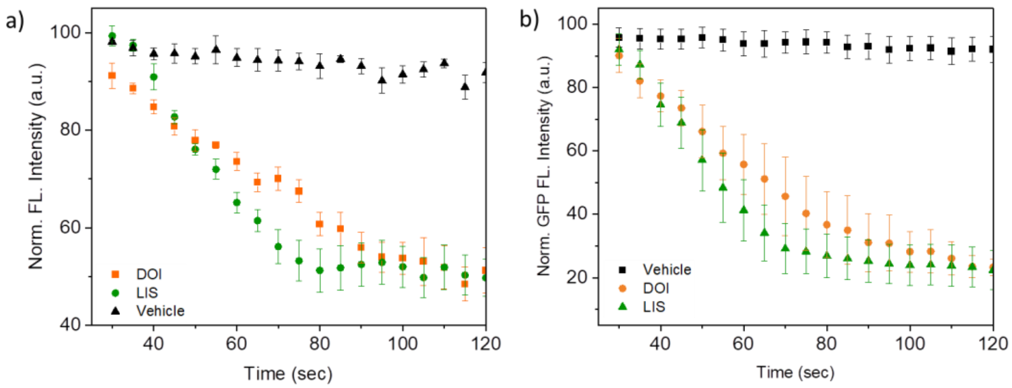


**Fig. S12**| Zoomed-in representation of 25−120 s time scale to show (a) PI(4,5)P2 depletion tracked by **MFR-12aa** (Fig. 3g) , and (b) GFP tagged 5-$HT_{2A}$ receptor internalization upon agonist (DOI and Lisuride) addition (Fig. S11e, bottom). Data represented as the mean ± SEM ($N$=3).

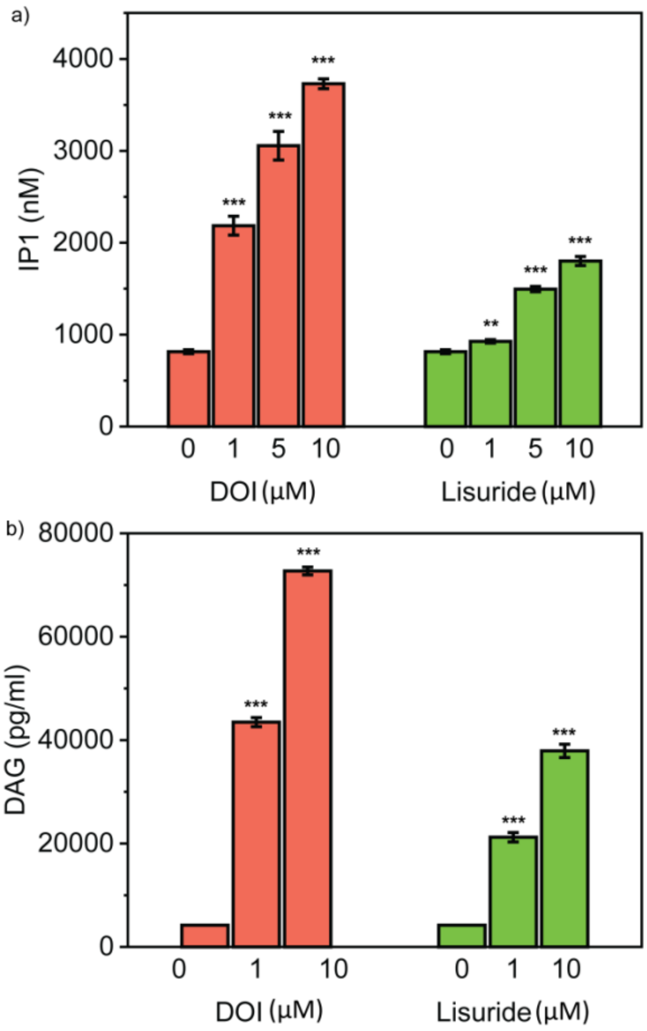


**Fig. S13|** IP1 and DAG assay upon DOI and Lisuride stimulation: HEK293 cells stably expressing human 5-$HT_{2A}$ receptor were either stimulated with increasing doses of DOI (hallucinogen) or Lisuride (non-hallucinogen), followed by measurement of intracellular (a) IP1 (nM) and (b) DAG (pg/mL) production using commercially available ELISA kits. Lisuride induced IP1 and DAG production was significantly lower at all the doses tested. Data has been expressed as the mean ± SEM of three independent experiments performed in duplicates. The significance of the differences was evaluated by one-way ANOVA followed by Tukey's post hoc test of the Software 'GraphPad Prism 5.0' (GraphPad Software, Inc., San Diego, CA, USA). The values of *$p < 0.05$ with respect to basal values were considered to be statistically significant.

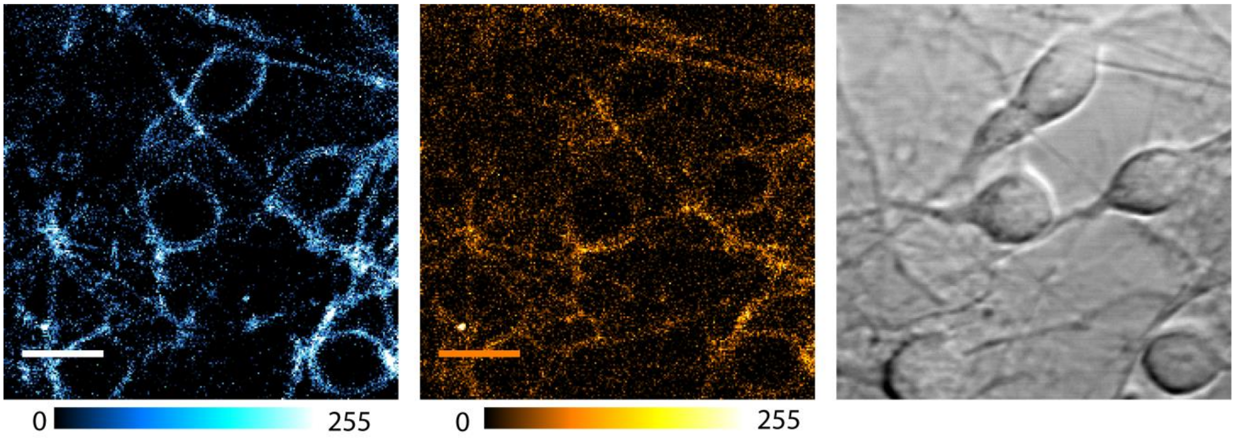


**Fig. S14|** Representative confocal single z-plane images of cortical neurons incubated with **MFR-12aa** sensor (1 µM, 5 min). **MFR-12aa** was excited with 780 nm, 2-photon laser. Fluorescence emission: blue channel ($\lambda_{em}$: 480−520 nm); orange channel ($\lambda_{em}$: 580−620 nm).

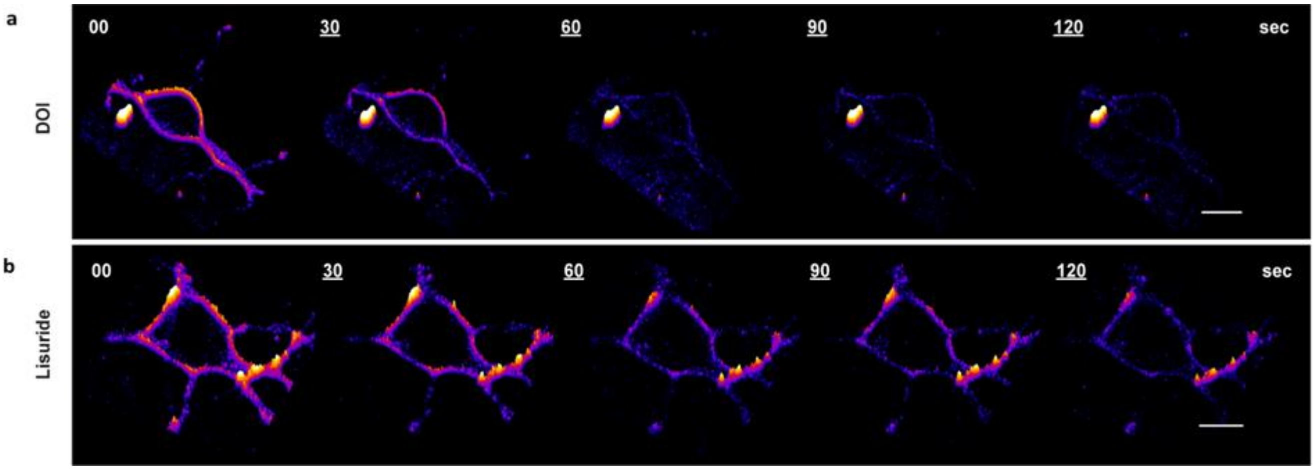


**Fig. S15|** Zoomed-in time frames (Fig. 4c, d) of confocal time-lapse imaging of rat primary cortical neurons incubated with **MFR-12aa** (1 µM), imaged at $\lambda_{ex}$ 780 nm, $\lambda_{em}$ 480-520 nm (blue channel), scale bar 10 µm. 3D intensity time traces for a single z-slice during ligand stimulation with (a) DOI (10 µM) and (b) Lisuride (10 µM).

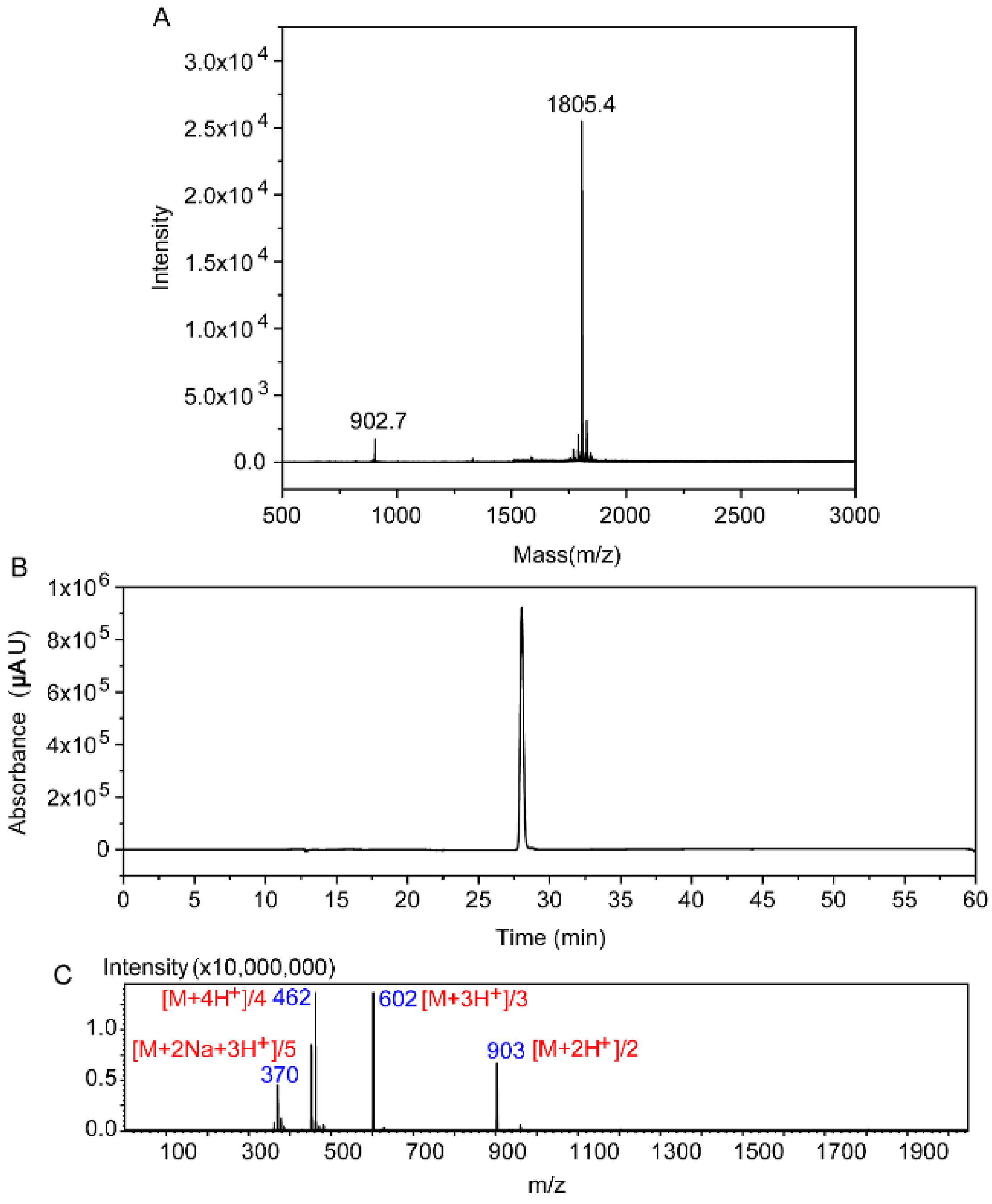


**Figure S16|** (a) MALDI-TOF MS spectra of **MFR-12aa** depicting the product peak at m/z 1805.4 (b) LC trace for **MFR-12aa** at 380 nm. (c) Positive mode ESI/MS depicting the product peak at different m/z values.

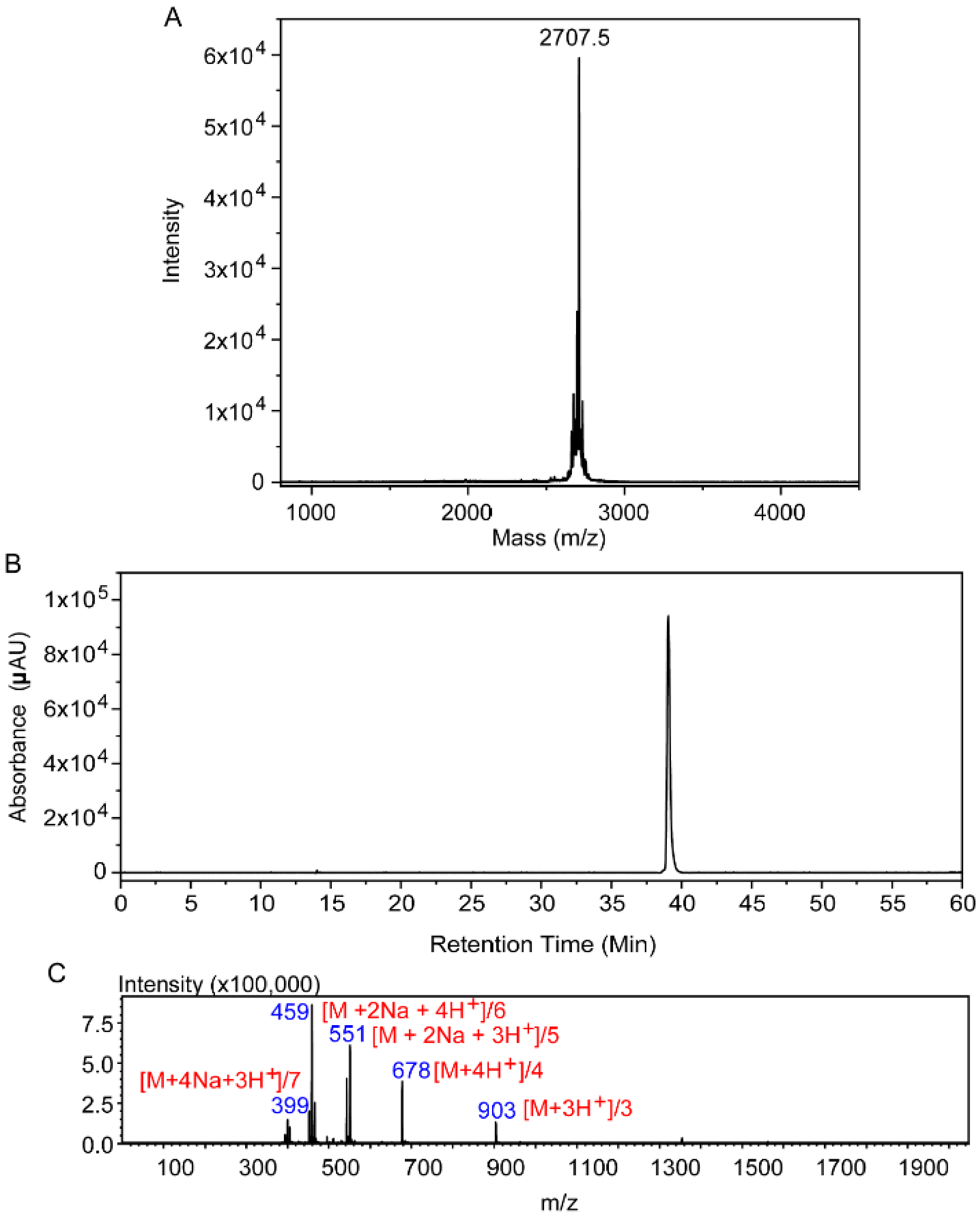


**Figure S17|** (a) MALDI-TOF MS spectra of **MFR-20aa** depicting the product peak at m/z 2707.5 (b) LC trace for **MFR-20aa** at 380 nm. (c) Positive mode ESI/MS depicting the product peak at different m/z values.

**Table S1. Previously reported fluorescent PI(4,5)P2 sensors.**

| Sensor | Response | Incorporation | Advantage | Existing Issues |
|---|---|---|---|---|
| **PLCδ-PH-GFP**[7] | Not responsive | Transfection/ electroporation | Expressible in cells | High background signal; non-quantitative response; cellular incorporation and expression vary from cell-to-cell; also binds to inositol-trisphosphate; diffusion limited slow response. |
| **TopFluor® PI(4,5)P2**[8] | Not responsive | Cell permeable | Cell permeable | Direct modification of lipid hence properties different from native lipid; Possibility of altered cellular localization and often cannot replicate biological function of native lipids. Irreversible response and hence cannot image dynamics. |
| **DAN–eENTH**[9] | Ratio metric | Micro-injection-based incorporation is limited to single-cell imaging. | Responsive and low background signal; Quantification possible | Tedious cellular incorporation possible in only one cell at a time; Imaging in tissues and multi-cellular organisms not straight forward and not reported; diffusion limited slow response. |
| **Lyn-PlcR (YFP-PH-PL-eCFP)**[10] | FRET based | Transfection | Quantification possible | Low dynamic range leading to low sensitivity; Tedious cellular incorporation; imaging in tissues and multi-cellular organisms not straightforward and not reported; diffusion limited slow response. |
| **DAN-13aa**[4] | Ratiometric peptide based | Cell permeable | Cell permeable and quantitative | Also responds to PI4P and dynamic tracking not reported. |

**Table. S2|** Summary of MD simulations performed for designing the PI(4,5)P2 selective probes.

| Simulations | Length | Bilayer composition |
|---|---|---|
| PI(4,5)P2-Gel-20aa | 200 ns*1, 30 ns*2 | POPC: PI(4,5)P2 |
| PI(4)P-Gel-20aa | 200 ns*1, 30 ns*2 | POPC: PI(4)P |
| PI(3,4,5)P3-Gel-20aa | 200 ns*1, 30 ns*2 | POPC: PI(3,4,5)P3 |
| PI(4,5)P2-Gel-12aa | 30 ns*3 | POPC: PI(4,5)P2 |
| PI(4)P- Gel-12aa | 30 ns*3 | POPC: PI(4)P |
| PI(3,4,5)P3- Gel-12aa | 30 ns*3 | POPC: PI(3,4,5)P3 |

**Table. S3|** Extent of maximum N-terminal dipping and average binding energy for each simulated system during 30 ns simulations.

| **Simulations** | **Extent of N-terminus dipping (Å)** | **Average binding energy (Kcal/mol)** |
|---|---|---|
| PI(4,5)P2- Gel-20aa | -14.4 | -282.2±32.9 |
| PI(4)P- Gel-20aa | -4.2 | -190.7±44.2 |
| PI(3,4,5)P3- Gel-20aa | -11.2 | -420.0±89.5 |
| PI(4,5)P2- Gel-12aa | -19.6 | -281.4±15.8 |
| PI(4)P- Gel-12aa | -6.2 | -147±17.5 |
| PI(3,4,5)P3- Gel-12aa | -17.0 | -306.7±31.7 |

Calculations were performed on two independent 30 ns and first 30 ns of the 200 ns run for Gel-20aa-phosphoinositide membrane systems and three independent 30 ns runs of Gel-12aa-phosphoinositide membrane systems.
