## Supplementary material for "A Cell-Permeable Fluorescent Probe Reveals Temporally Diverse PI(4,5)P2 Dynamics Evoked by Distinct GPCR Agonists in Neurons": video-description

**Video-1: Representative molecular dynamics simulations of PI(4,5)P2, PI(4)P, and PI(3,4,5)P3-POPC membranes with Gel-20aa binding peptide.** Simulations showing Gel-20aa-PIP interactions (PIP gray licorice, phosphate O-atoms in red) in a POPC (grey lines) membrane bilayer. Peptide is shown in purple (new cartoon) and cationic sidechains on the peptide are shown in blue (licorice, Arg) and green (licorice, Lys). Video segments, PI(4,5)P2 (left), PI(4)P (middle) and PI(3,4,5)P3 (right), are taken around the highest dipping position of the peptide N-terminus for 15 ns of each 200 ns trajectories.

**Video-2: Representative molecular dynamics simulations of PI(4,5)P2, PI(4)P, and PI(3,4,5)P3-POPC-membranes with Gel-12aa peptide.** Simulations depict Gel-12aa-PIP interactions, with PIP represented in gray licorice and phosphate O-atoms in red, within a POPC membrane bilayer (depicted by grey lines). The peptide is illustrated in purple (new cartoon), with cationic sidechains shown in blue (Arg), green (Lys), and orange (Val) licorice. Each video segment, PI(4,5)P2 (left), PI(4)P (middle), and PI(3,4,5)P3 (right), displays a representative 30 ns trajectory.

**Video-3: 360° view image from confocal images recorded at the head region of a PIP5K overexpressing transgenic (*gqIs25*) worm.** Worms were incubated with MFR-12aa (10 μM) for 1 h.

**Video-4: 360° view image from confocal images recorded at the head region of a wild-type (*N2*) worm.** Worms were incubated with MFR-12aa (10 μM) for 1 h.

**Video-5: PI(4,5)$P_2$ dynamics in live $5HT_{2A}$ over-expressing HEK293T cells treated with 5-HT and control experiment with Wortmannin.** Confocal time lapse imaging depicting substrate dependent PI(4,5)P2 hydrolysis and replenishment in cells incubated with MFR-12aa (5 μM); two-photon excitation $\lambda_{ex}$ 780 nm, $\lambda_{em}$ blue channel (480-520 nm), scale bar 10 μm. Left video shows PI(4,5)P2 depletion during stimulation with 5-HT (10 μM) for a single z-slice. Data was acquired for unstimulated cells (1 min), substrate stimulation (3 min), and substrate removal and replacement with fresh media (3 min). Right video shows cells treated with Wortmannin (10 μM) under identical experimental conditions. 3D intensity scale representation (0-255).

**Video-6: Buffer control and serotonin-wortmannin induced PI(4,5)$P_2$ dynamics in live $5HT_{2A}$ over-expressing HEK293T cells.** Confocal time lapse imaging depicting substrate dependent PI(4,5)P2 hydrolysis and replenishment in cells incubated with MFR-12aa (5 μM); two-photon excitation $\lambda_{ex}$ 780 nm, $\lambda_{em}$ blue channel (480-520 nm), scale bar 10 μm. Video trace for a single z-slice, 3D intensity scale representation (0-255) during addition of Buffer (left) and stimulation with 5-HT and Wortmannin (10 μM each) (right), indicated in the respective time frames. Data was acquired for unstimulated cells (1 min), substrate stimulation (3 min), and substrate removal and replacement with fresh media (3 min).

**Video-7: DOI and Lisuride induced PI(4,5)$P_2$ dynamics in live $5HT_{2A}$ over-expressing HEK293T cells.** Confocal time lapse imaging depicting substrate dependent PI(4,5)P2 hydrolysis and replenishment in cells incubated with MFR-12aa (5 μM); two-photon excitation $\lambda_{ex}$ 780 nm, $\lambda_{em}$ blue channel (480-520 nm), scale bar 10 μm. Video trace for a single z-slice, 3D intensity scale representation (0-255) during stimulation of DOI (10 μM) (left) and Lisuride (10 μM) (right) indicated in the respective time frames. Data was acquired for unstimulated cells (1 min), substrate stimulation (3 min), and substrate removal and replacement with fresh media (3 min).

**Video-8: Buffer control and Serotonin induced GFP-tagged $5HT_{2A}$ receptor dynamics in live $5HT_{2A}$ over-expressing HEK293T cells.** Confocal time lapse imaging depicting substrate dependent $5HT_{2A}$ receptor internalization by tracking GFP fluorophore; $\lambda_{ex}$ 488 nm, $\lambda_{em}$ blue channel (500-560 nm), scale bar 10 µm. Video trace for a single z-slice, 3D intensity scale representation (0-255) during addition of buffer (left) and stimulation with 5-HT (10 µM) (right), indicated in the respective time frames. Data was acquired for unstimulated cells (1 min), substrate stimulation (3 min) and substrate removal and replacement with fresh media (3 min).

**Video-9: DOI and Lisuride induced GFP-tagged $5HT_{2A}$ receptor dynamics in live $5HT_{2A}$ over-expressing HEK293T cells**. Confocal time lapse imaging depicting substrate dependent $5HT_{2A}$ receptor internalization by tracking GFP fluorophore; $\lambda_{ex}$ 488 nm, $\lambda_{em}$ blue channel (500-560 nm), scale bar 10 µm. Video trace for a single z-slice, 3D intensity scale representation (0-255) during stimulation of DOI (10 µM) (left) and Lisuride (10 µM) (right) indicated in the respective time frames. Data was acquired for unstimulated cells (1 min), substrate stimulation (3 min) and substrate removal and replacement with fresh media (3 min).

**Video-10: Buffer control and serotonin induced $PI(4,5)P_2$ localized dynamics in cortical neurons.** Confocal time lapse imaging depicting substrate dependent PI(4,5)P2 hydrolysis in neurons incubated with MFR-12aa (1 µM); two-photon excitation $\lambda_{ex}$ 780 nm, $\lambda_{em}$ blue channel (480-520 nm), scale bar 10 µm. Video trace for a single z-slice, 3D intensity scale representation (0-255) during addition of buffer (left) and stimulation with 5-HT (10 µM) (right), indicated in the respective time frames. Data was acquired for unstimulated cells (1 min), substrate stimulation (3 min) and substrate removal and replacement with fresh media (3 min).

**Video-11: DOI and Lisuride induced $PI(4,5)P_2$ localized dynamics in cortical neurons.** Confocal time lapse imaging depicting substrate dependent PI(4,5)P2 hydrolysis in neurons incubated with MFR-12aa (1 µM); two-photon excitation $\lambda_{ex}$ 780 nm, $\lambda_{em}$ blue channel (480-520 nm), scale bar 10 µm. Video trace for a single z-slice, 3D intensity scale representation (0-255) during stimulation of DOI (10 µM) (left) and Lisuride (10 µM) (right) indicated in the respective time frames. Data was acquired for unstimulated cells (1 min), substrate stimulation (3 min) and substrate removal and replacement with fresh media (3 min).
